# Chromosomal inversions assist acute salinity and temperature adaptation in Atlantic cod (*Gadus Morhua*) eggs

**DOI:** 10.1101/2025.09.27.678988

**Authors:** Rebecca Krohman, Simon Henriksson, Esben Moland Olsen, Halvor Knutsen, Rebekah A. Oomen

## Abstract

Chromosomal inversions, on chromosomes 2, 7, and 12 in Atlantic cod (*Gadus morhua*), have been consistently associated with environmental gradients, particularly salinity and temperature, across the species’ natural range. While these correlations suggest a role for local adaptation, the underlying selective mechanisms remain unresolved. Critically, few experiments have yet tested the fitness consequences of these inversions under controlled environmental conditions. Such work is essential to establish causality. Here, we conducted an acute stressor experiment with ten treatments of combined temperature and low salinity stress. Cod eggs (N = 1265) were placed into each treatment for 24 hours, and the number of individuals that floated initially and finally was counted. The ability of an egg to float is directly related to its survival, as sunk individuals cannot survive, and individuals lose buoyancy after they die. Inversion genotypes for the inversions on chromosomes 2, 7, and 12 were determined for individuals from the T03-S25, T07-S25, T16-S25, and T16-S35 treatments. We found that sinking increased with temperature and that more individuals sank in 25 ppt than in 35 ppt after 24 hours. Interestingly, some individuals in the 25 ppt treatments recovered (sank initially but floated finally) after 24 hours. We found evidence that the inversions on chromosomes 2 and 12 might assist with temperature and salinity adaptation, respectively, in cod eggs, while the inversion on chromosome 7 did not affect floating. Our results can inform the management and conservation of this endangered species.

**Summary statement:** We show that chromosomal inversions found in Atlantic cod help with adaptation to short-term temperature and salinity stress at the egg stage.

## 1 Introduction

A stressor is any biotic or abiotic variable that exceeds the normal range to which an animal or population is adapted, resulting in physiological stress or demographic decline (Barrett et al., 1976). For example, an individual can adapt to water temperatures up to 25°C, but an extreme heat event can increase temperatures to 30°C, causing the individual physiological stress. Laboratory-based stressor experiments allow for rigorous control of confounding variables, allowing for strong conclusions to be made about the response to stressors (Nielsen et al., 2009). Experiments can be informative to management and conservation about the ecosystems, species, and populations most at risk due to climate change (Côté et al., 2016). However, most experimental designs rely on discrete, binary treatments (e.g., temperatures at 10°C and 20°C) rather than on a continuous scale (ex., temperatures at 10,12, 14, 16, 18, and 20°C). Experiments with multiple levels of a given stressor build stronger models to predict responses across a range of stress conditions (Orr et al., 2024). Furthermore, there are few stressor studies which take intraspecific variation into account, but these studies can provide information on which populations are at the highest risk (Nagelkerken et al., 2023; Orr et al., 2020). Overall, understanding genetically based variation in response to ecologically relevant stressors at continuous scales will fill a knowledge gap (Nagelkerken et al., 2023; Côté et al., 2016).

Reductions in ocean salinity, particularly below 34 ppt, occur naturally (Fransson et al., 2015) but are increasingly driven by climate change. Enhanced precipitation, elevated temperatures, and earlier seasonal freshwater runoff are leading to greater and more variable freshwater input into coastal and shelf ecosystems (Westra et al., 2014; Cyr et al., 2021; 27). Teleosts can regulate internal salinity levels; however, increased regulation of internal conditions impacts energy demands (Lambert et al., 1994), and early life stages do not regulate internal conditions well (Ackerly et al., 2023).

The co-occurrence of temperature and low salinity stress is common in the marine system. Due to increasing temperatures, glacier melt in the Arctic is high, and ocean currents bring cold freshwater into Northern Atlantic marine ecosystems (Cyr et al., 2021). Northern areas of Norway, including fjords that are naturally fresh, experience warming due to the lack of glaciers providing cool waters (Tesdal et al., 2020). Temperature stress tends to have a stronger impact than salinity, and salinity interacts with temperature to increase negative responses (Magill and Sayer, 2004; Bian et al., 2016). Most studies on the impact of temperature and salinity stress in fish focus on biochemical physiology (Magill and Sayer, 2004; Árnason et al., 2013; Dutil et al., 1992) and there is a lack of studies focusing on population-level effects.

Maternal effects are the epigenetic cues and provisions passed from a mother onto her offspring (Wolf and Wade, 2009). When an individual experiences stress, epigenetic changes can occur and be passed onto offspring, allowing the offspring to be better adapted to the stress (Youngson and Whitelaw, 2008). Therefore, maternal effects are a large confounding variable in stressor studies, increasing data set variation (Youngson and Whitelaw, 2008; Vega-Trejo et al., 2018). For example, the conditions that cod mothers are exposed to during spawning impact the survival, buoyancy, and hatching of cod eggs (Schmidt et al., 2024). However, maternal effects are rarely taken into account in studies focusing on fish early life stages (Catalán et al., 2020). Generally, experimental studies control for maternal effects by experimenting on an F2 generation. However, the lack of F2 generations in non-model organisms with long generation times inhibits accounting for maternal effects.

Cod is an endangered species (Sobel, 1996) due to overfishing (Hamilton et al., 2004). Cod have had difficulty rebounding (DFO, 2023), possibly due to environmental stressors (Hilborn and Litzinger, 2009). Cod spawns in North Atlantic rocky shores and banks (Zemeckis et al., 2014) in temperatures between 5 and 10°C (Righton et al., 2010) and salinities down to 15 ppt (Nissling and Westin, 1997). Cod eggs and larvae are neutrally buoyant and live in the upper levels of the water column (Nissling and Westin, 1997). A female can spawn millions of eggs across a spawning season, but few will make it to maturity (Kjesbu et al., 1998). Therefore, there is high mortality of cod during early life and early life stages are more sensitive to stressors (Hasan et al., 2023).

Temperature impacts cod growth rates (Otterlei et al., 1999; Laurence and Rogers, 1976) while salinity impacts buoyancy of eggs, larvae, and juveniles (Schmidt et al., 2024). The extent to which temperature impacts larval development rate is population dependent and likely due to genetic differences (Oomen and Hutchings, 2015). However, one estimate suggests that at 7°C individuals hatch 20 days post-fertilization (Pepin et al., 1997). Temperature has a larger impact on the survival of larval stages, while salinity impacts egg stage mortality (Jordaan and Kling, 2003; Schmidt et al., 2024). Furthermore, stress responses, including mortality and feeding rates of larval cod, increase at the extremes of the normal spawning ranges for temperature and salinity (Laurence and Rogers, 1976; Jordaan and Kling, 2003). Salinity and temperature have an interactive effect on adaptation (Laurence and Rogers, 1976) and might be seasonally dependent (Magill and Sayer, 2004).

Local adaptation in cod has been partly attributed to chromosomal inversions. Inversions are a flipped part of the genome that evolves separately from the rest of the genome due to recombination suppression in heterozygotes (Kirkpatrick, 2010; Wellenreuther and Bernatchez, 2018). Therefore, inversions can speed up evolutionary processes and might be important for adaptation to climate change (Oomen et al., 2020). Inversions have been correlated with environmental conditions in many organisms (*Pecten maximus* and temperature [Hollenbeck et al., 2022], *Zea mays* and altitude [Fang et al., 2012], *Coelopa frigida* and salinity [Mérot et al., 2021]), including teleost species (*Mallotus villosus* and temperature [Cayuela et al., 2019]). Discovering the relationships between environmental variables and inversion frequencies is an increasing research area, with many studies being observational. However, experiments are needed to understand the mechanism underlying the relationships between inversion frequencies and environmental variables (Hoffmann et al., 2004; Wellenreuther et al., 2025). Common garden experiments, which control environmental conditions, can reveal the causal adaptive impacts of inversions by removing potential confounding variables (Mérot et al., 2020; Whitlock, 2015).

Three major inversions in the cod genome are correlated with environmental variables on chromosomes 2, 7, and 12 (hereafter, the inversions are referred to as inv2, inv7, and inv12, respectively). For example, inv2 and inv12 are strongly correlated with low salinity and temperatures, respectively (Bradbury et al., 2010; Berg et al., 2015; Link et al., 2009). Environmental correlations with inv7 are less clear, but trends have been found with salinity, oxygen, and temperature (Sodeland et al., 2016; Bradbury et al., 2010; Sinclair-Waters et al., 2018; Hemmer-Hansen et al., 2013). The inversions likely interact with each other as well, specifically inv7 with inv2 and inv12 (Sodeland et al, 2022; Oomen, 2019). However, there is little understanding of the role of inversions in adaptation to temperature and salinity simultaneously.

We conducted controlled stressor experiments to understand how Atlantic cod eggs respond to acute temperature and salinity stress. The experimental design included four temperatures (3, 7, 12, and 16 °C) and two salinity levels (25 and 35 ppt). We predicted that sinking would be higher in the thermal extremes due to individuals being under higher stress and losing buoyancy. We also expected temperature and salinity to interact, and there to be more sinking when 3 °C and 16°C were applied with 25 ppt than when applied separately. We tested the hypothesis that the genotypes of the individuals floating after 24 hours would be dependent on the treatment. For example, individuals with the inv2 derived homozygote or heterozygote genotypes might be more likely to float in treatments with a salinity of 25 ppt, while salinity will not impact inv 7 and 12 frequencies. We predicted a positive association between temperature and the frequency of derived homozygotes at inv7 and a negative association at inv12, while expecting no temperature effect on inv2 genotype frequencies.

## 2 Materials and Methods

All salinities were measured using a YSI ProSolo meter with an ODO/CT probe (±1.0% of reading), temperatures were measured using a Topfin Digital Thermometer (±1°C) or a TEDDINGTON TL310 digital thermometer (±1°C), and all pH, nitrates, nitrites, and ammonia were measured using an API saltwater master test kit, unless otherwise stated. Freshwater was collected from a Synergy UV-R system filter and was cooled to ambient temperatures. All saltwater was collected from 75 m deep offshore and macrofiltered with a 40 μm filter before flowing into lab tanks. Individuals were held in one of three circular plastic containers with a 200 μm mesh bottom and a 19 cm inside edge diameter in a holding tank with low water surface movement before experimentation (Fig. 1). During experimentation, this water’s average (±SE) temperature, salinity, and pH were 7.1 (±0.29)°C, 33.7 (±1.10) ppt, and 8.0 (±0.03), respectively. Lighting in the laboratory during all experimentation was on a 10-hour light, 14-hour dark schedule. Measured lux in the stressor experimentation area was 503 and 210 in the holding tank area (measured using a Biltema light meter Art 15-341). Air temperatures within the laboratory during experimentation ranged between 3°C and 10.5°C but generally stayed around 8°C. In the laboratory, eggs were moved, collected, and placed using a plastic pipette with the end cut off, approximately 4 mm in diameter. All analyses used to classify SNP data and run statistics were conducted in R using R Studio v4.4.2 (R Core Team, 2023). Figures were created in GraphPad Prism v10.4.1 or using the ggplot2 package in R (v3.5.2; Wickham, 2016). An alpha value of 0.05 was used throughout. All model outputs were predicted with the ggpredict function in the ggeffects package (v2.2.1; Lüdecke, 2018) for graphing purposes. We had approval from the Animal Ethical Committee of the Norwegian Food Safety Authority (Mattilsynet) to carry out the work with the utmost care for environmental impacts and fish welfare (FOTS ID 28408).

**Fig. 1.**
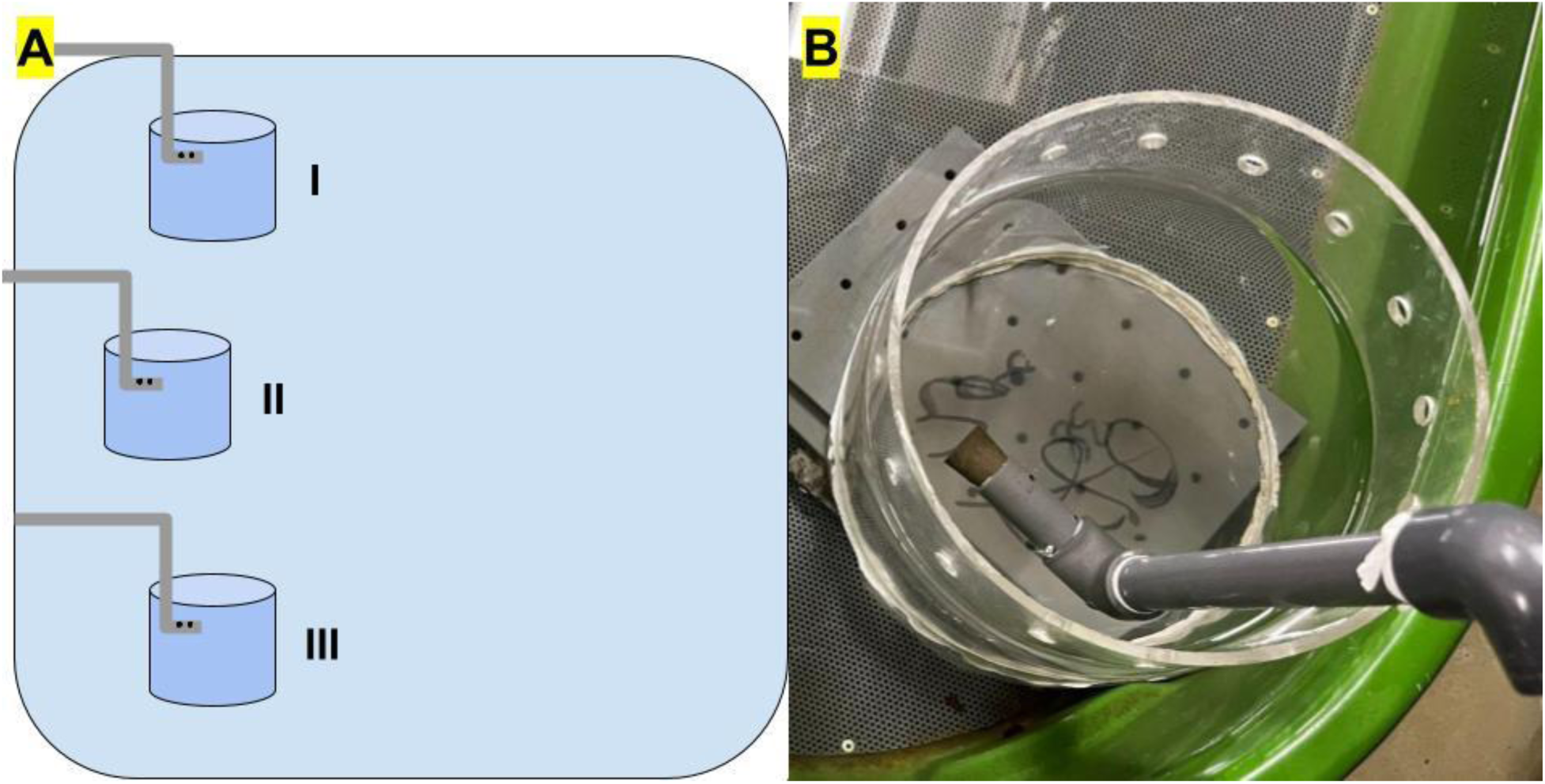
A schematic of the setup of the holding tank and holding containers I, II, and III, in which individuals were held before experimentation (A) and a close-up of one of the holding tanks (B). The schematic shows the tubing that the water flowed through (grey), the bore holes at the end of the tubing that helped control water flow (black), and the small circular holding containers (dark blue) inside the large holding tank (light blue). Note the mesh at the bottom of the holding container in part B. The grey blocks below the holding container were used to prop the holding container higher so that the holes on the top of the container would be above the water line to prevent any eggs from escaping through the holes.

### 2.1 Spawning population

Along the Skagerrak coast, Atlantic cod (Linnaeus, 1758) are polymorphic for LG2, LG7, and LG12. We collected 45 (hereafter F0) individuals from two sampling sites in Norway (N=45: 17 from approximately 58.4° latitude 8.8° longitude; 28 from approximately 58.3° latitude and 8.6° longitude) for spawning in a controlled large, outdoor, semi-natural human-made pond with a diameter of 50 m and a maximum depth of 4 m. The pond has flow-through seawater from 75m deep and is located at the Flødevigen Research Station, Institute of Marine Research in Norway (Fig. 2). The F0 individuals spawned freely to create the F1 generation in Winter 2022 and then were removed from the pond and released back into the wild. Approximately 50 individuals from the F1 generation grew until maturity and spawned freely to produce the F2 generation in winter 2024.

**Fig. 2.**
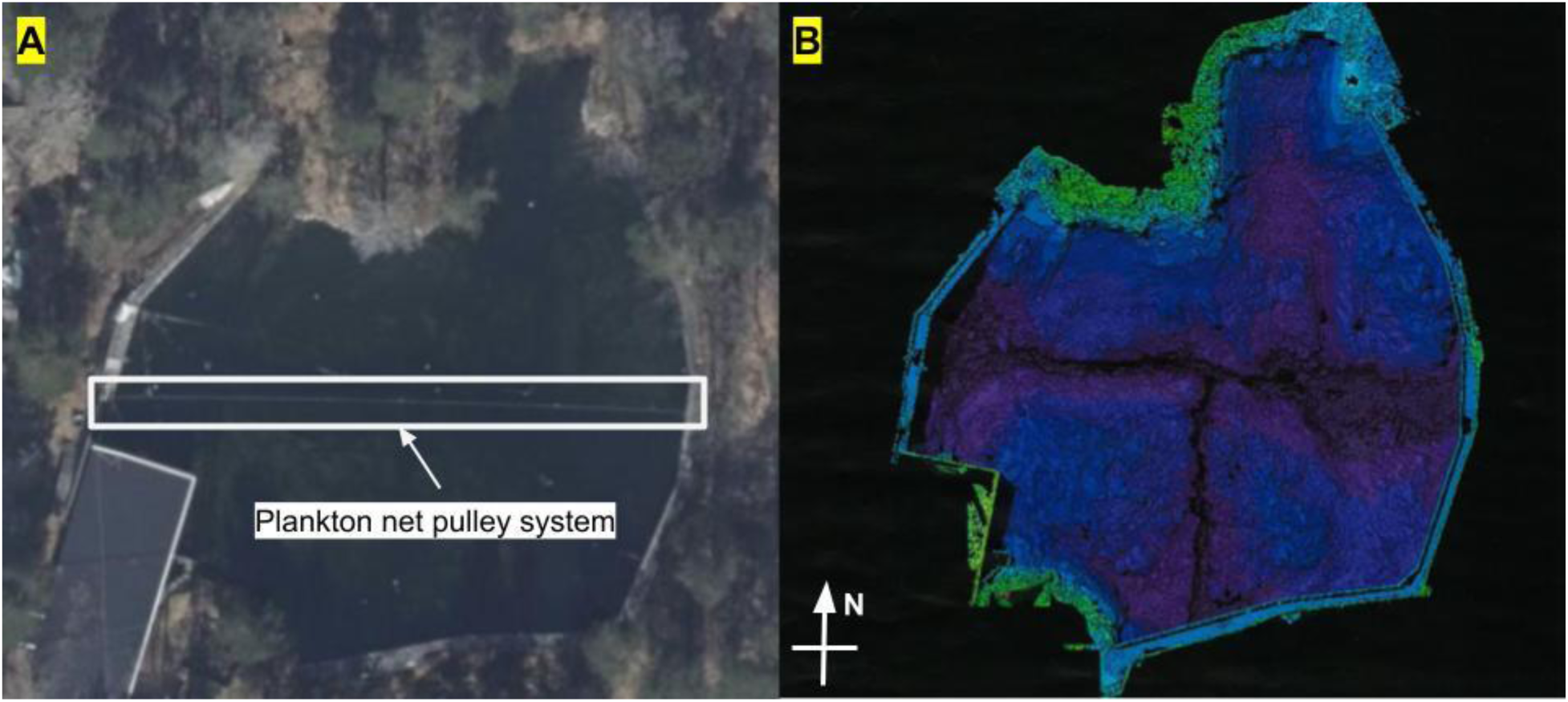
Satellite photo of the cod pond from 222 (A) and depth scan of the cod pond (B). The plankton net pulley system set up across the pond is marked in white.

### 2.2 Collection and holding

Eggs were collected by pulling a plankton net across the pond using a pulley system. The plankton net drags across the pond’s surface along the deepest part of the pond (Fig. 2). The plankton net, with a mesh size of 320 μm and a diameter of 46 cm, is connected by rope to a buoy on the pulley system and sits about 1.5m away from the buoy. We collected 4 batches of eggs on February 10th, 17th, and 25th, and March 3rd, 2025, by pulling the plankton net across the pond twice. We pulled the plankton net 10 m min-1 in one direction and 16.7 m min-1 back in the opposite direction to collect individuals at and just below the surface. Eggs were immediately placed into a bucket with water collected from the surface of the pond. The pond water conditions were measured 1 meter deep during collection using a YSI ProSolo meter with an ODO/CT probe, with the average (±SE) temperature, salinity, and dissolved oxygen being 4.2 (±0.69)°C, 33.49 (±0.15)%, and 98.1 (±9.09)%, respectively. The average (±SE) pH, ammonia, nitrate, and nitrite of the pond surface water were measured to be 7.975 (±0.03), 0.04 (±0.04), 0 (±0), and 0 (±0), respectively. The eggs were then brought into the lab and sorted, using a Leica ES2 dissecting microscope, into early stage (body length < 50% of the egg perimeter; stages 1 and 2) and late stage (body length > 50% of the egg perimeter; stages 3-5, following Thompson and Riley, 1981). Eggs were separated by stage and placed into holding containers (Fig. 1). Batches 1 and 2 were collected as stated, but there were some small changes to the collection of batches 3 and 4 (resulting in the presence of batch 4b) that can be found in Appendix B.

### 2.3 Experimental protocol

Ten treatments (determined from pilot experiments, see Appendix A) were used for the temperature and salinity stressor experiment: T stands for temperature in °C and S stands for salinity in ppt, T03-S25, T03-S35, T07-S20, T07-S25, T03-S30, T07-S35, T12-S25, T12-S35, T16-S25, and T16-S35. The T07-S35 treatment was the control, while the T12 treatments acted as a positive temperature control. The T07-S20 and T07-S30 treatments were negative and positive salinity controls, respectively. Lowered salinity was created by mixing fresh water with a pH of 8.0 (raised to 8.0 by adding API pH up for consistent pHs across treatments) with salt water. The salinity, pH, ammonia, nitrate, and nitrite were measured before treatment to make sure they were within safe ranges (7.9-8.0 pH, >0.5 ammonia, >0.1 nitrate, >0.1 nitrite). Temperature was controlled in the treatments using a thermal gradient block (Appendix B; Veenhof et al., 2024). At least one hour before the start of a replicate, 200 mL of treatment water was placed in a 250 mL Borosilicate glass 3.3 beaker, and the beaker was placed in the respective column of the thermal gradient block (column A for 16°C, D for 12°C, H for 7°C, and K for 3°C; Fig. 3). In this hour, the beaker temperature rose or lowered to its respective temperature.

**Fig. 3.**
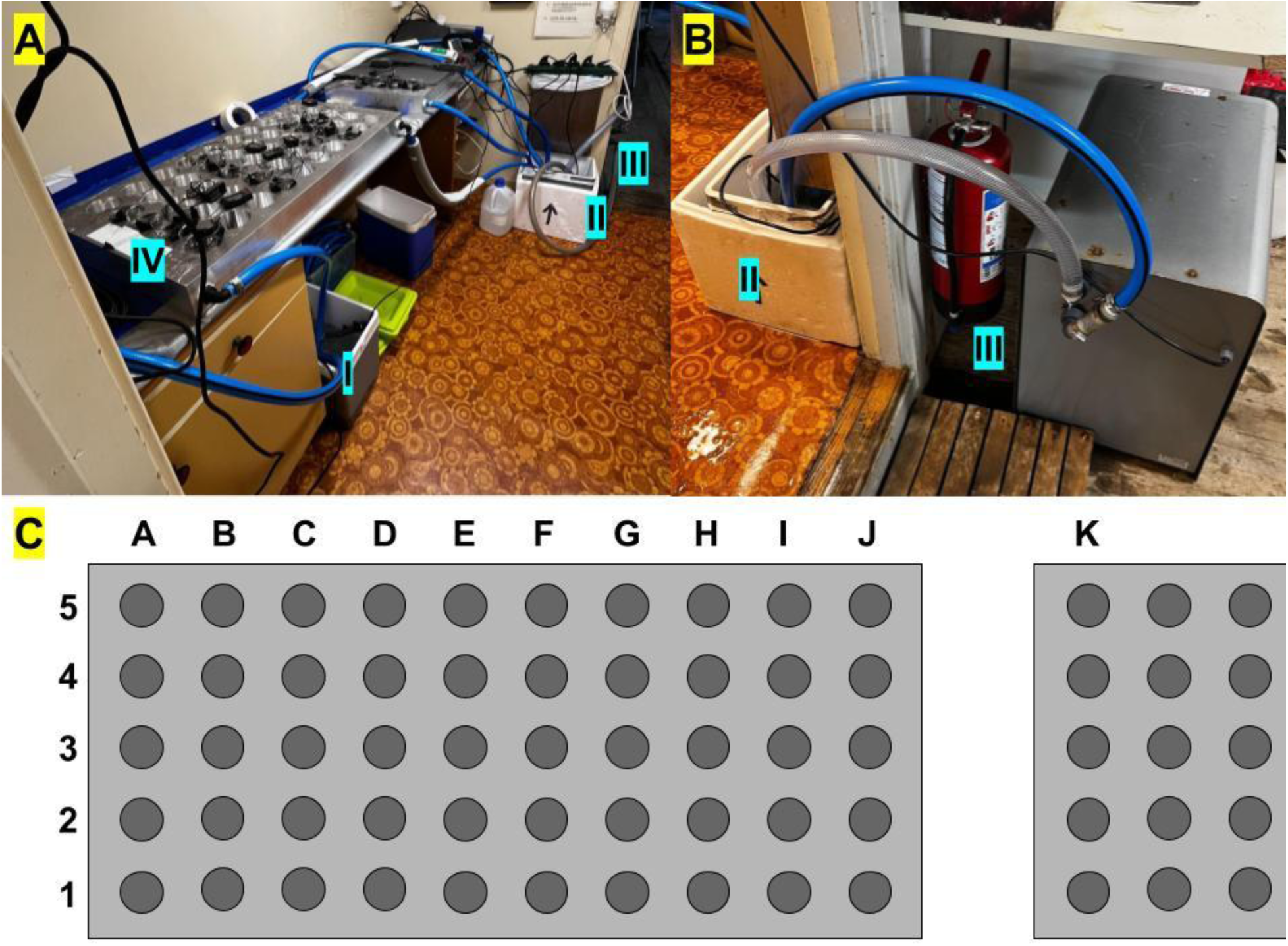
The setup of the thermal gradient plates used for the temperature salinity stressor experiment (A), including the “warm” cooler (I), “cool” cooler (II), chiller (III), and temperature controller (IV). A close look at the connection between the “cool” cooler (II) and the chiller (III; B). A schematic of the two temperature gradient blocks used in experimentation with labels for rows 1 to 5 and columns A to K (C).

In a given replicate, 10 individuals with a known stage were placed into each treatment, 5 minutes apart. All eggs were held in the holding tank for at least 20.25 hours before experimentation, and all experiments were started within 74.08 hours of collection. 15 minutes after individuals were placed in the treatments, the number of individuals floating and sunk in each stage was counted. The number of individuals floating was used as a proxy for individuals alive, while the number of individuals sunk was used as a proxy for individuals dead. 12 and 24 hours after individuals were placed in the first treatment in a given replicate, each beaker was stirred slowly twice in 5-second intervals. After 24 hours of treatment, the number of individuals floating and sunk in the beaker was counted, and individuals were staged visually using a Leica ES2 dissecting microscope when needed. The temperature of each beaker was measured at 0, 12, and 24 hours. All individuals in the T03-S25, T03-S35, T07-S25, T16-S25, and T16-S35 were sampled for DNA extraction. 13 replicates were completed for treatments T03-S25, T03-S35, T07-S25, T12-S25, T12-S35, T16-S25, and T16-S35, while 12 replicates were completed for treatments T07-S20, T07-S30, and T07-S35. A total of 1265 individuals were used in the temperature and salinity stressor experiment (Table S1).

### 2.4 Stage changes classification

To understand how individuals changed stages while in treatment in each beaker, we created five classifications for salinity 25 treatments, early floated, sunk developed, developed floated, float developed, and late floated, and three classifications for salinity 35 treatments, early sank, developed, and sank. These classifications were created based on what a given egg’s initial and final stages and states (Table S2). The number of individuals in a given beaker was classified in a given stage change by comparing the early sunk difference, the early float difference, the late sunk difference, and the late float difference. The assumption that the least number of individuals as possible would change stage and state, and that changes only occurred once in one direction, was made throughout.

### 2.5 DNA preparation, extraction, and sequencing

Eggs taken for DNA extraction were fixed in 96% ethanol and held in the lab for a maximum of 36 hours before being placed in a freezer at -20°C. All eggs in the T16-S35 (n=129) treatment and all floating individuals in the T03-S25 (n=26), T07-S25 (n=31), and T16-S25 (n=33) treatments were shipped overnight to Identigen in Ireland for DNA extraction and genotyping (Shapero, 2013). We used a panel of 4000 SNP loci designed for genetic monitoring of Swedish cod populations (Henriksson et al., in prep.; *cf.* Andersson et al., 2024). We received SNP data for 381 individuals and 3561 loci. SNP data were also received for extra individuals from batch 2 (n=47), 3 (n=58), 4 (n=27), and 4b (n=33) that did not undergo treatment to better estimate batch genomic traits. Details regarding the assignment of inversion genotypes and ecotypes to the eggs can be found in Appendix B.

### 2.6 Statistical methods

Due to genomic data only being collected from floating individuals in treatments with a salinity of 25, stressor genetic results were compared to batch genetic frequencies (calculated from all T16S35 individuals plus extra individuals that did not undergo treatment) to understand the effect of the treatments. The glmmTMB function in the glmmTMB package (v1.1.11; Brooks et al., 2017) was used to build multiple models to test the effects of temperature and salinity on the response variables of interest from experimentation. In all models, temperature (numeric) and salinity (factor with 2 levels) were interactive explanatory variables, and beaker number (factor with 13 levels) nested in batch (factor with 4 levels), and time since collection (numeric) were random factors. In all models, 35 ppt salinity was the reference point. To correct for multiple models, the p-value outputs of all models underwent false discovery rate correction. First, the response variables of the number of individuals floating initially, finally, and the float difference (integers) were tested (Equation 1). Next, allele A frequency differences (beaker allele frequency subtracted from the batch allele frequency) for inv2, inv7, and inv12 were tested (Equation 3). For the inv2 model, batch was dropped as a random effect due to it creating errors in the model. Finally, a PCA was created with genotype frequency differences (a beaker genotype frequency subtracted from the batch genotype frequency) for each beaker to account for multiple possible interacting response variables. The location of each beaker along the first, second, and third PCA axes was extracted and acted as a response variable representing genotype frequency differences (Equation 4).

Two models (one for S25 and one for S35) were built using the glmmTMB function in the glmmTMB package (v1.1.11; Brooks et al., 2017) to understand the effect of temperature on stage changes (8 total models were made, one for each stage change). The response variable was the number of individuals undergoing the given stage change (integer), the explanatory variable was temperature (numeric) and beaker number (factor) nested in batch (factor) and time since collection (numeric) were included as random factors (Equation 2). To understand how ecotype impacted the number of individuals floating in each treatment, the distribution of the probability of belonging to the offshore ecotype was compared within each treatment to the overall batch distribution using a Kolmogorov-Smirnov test via the ks.test function in the dgof package (v1.5.1; Conover, 1972).

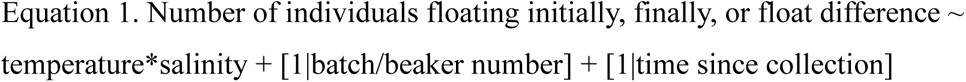

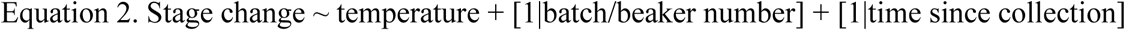

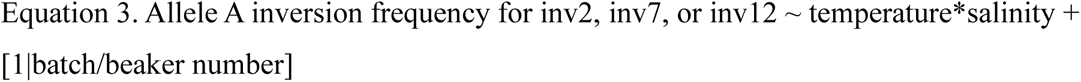

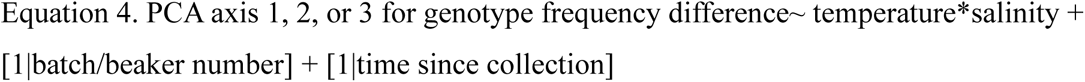

## 3 Results

### 3.1 Inversion and ecotype assignment results

The maximum and average±SE percent likelihood that the eggs belonged to the eastern Baltic cod ecotype was 0.01925 and 0.005903±0.000167, respectively, indicating that all eggs likely belonged to the coastal cod or offshore cod ecotypes. In all the batches, the heterozygote (0.417) and derived (0.574) genotypes for inv2 had about equal frequency, and the ancestral genotype had a low frequency (0.008; Fig. S1). The ancestral frequency was high (0.885) for inv7 compared to the heterozygote frequency (0.115; Fig. S1). Inv12 had a high frequency of derived genotypes (0.796) and a low frequency of heterozygotes (0.204; Fig. S1). Derived and ancestral genotypes were not present for inv7 and inv12, respectively. Alleles were labelled as ancestral allele and derived allele. Allele frequencies were calculated based on genotype frequencies. The frequency of the ancestral allele was 0.217, 0.945, and 0.102 across all batches for inv2, inv7, and inv12, respectively (Fig. S1).

### 3.2 The impact of temperature and salinity on cod eggs

In the T07-S20 treatment, we saw that all individuals sank initially and stayed sunk for the duration of the experiment. Individuals displayed both sinking and floating after 24 hours in the T07-S30 treatment, as some replicates had a negative float difference and some had a positive float difference (Fig. S2), but overall, 49% of individuals floated initially, and 66% floated finally. The number of individuals floating initially and finally in salinity 35 ppt was significantly different than salinity 25 ppt (Table 1). There was no effect of temperature on the number of individuals floating initially or finally (Table 1). Overall, in S25 treatments, 13% of individuals floated initially (14%, 16%, 11%, 12% at 3°C, 7°C, 12°C, and 16 °C, respectively), while all individuals floated initially in S35 treatments (Table 1; Fig. 4). After 24 hours, 77% and 96% of individuals floated in all S25 (80%, 75%, 84%, 73% at 3°C, 7°C, 12°C, and 16 °C, respectively) and S35 (99.2%, 99.2%, 97%, 90% at 3, 7, 12, and 16 °C, respectively) treatments, respectively (Table 1; Fig. 4). In the control treatment, T07S35, and in T03-S35, only one individual sank after 24 hours among all replicates. The calculated float difference was not significantly different between salinities 25 ppt and 35 ppt (Table 1; Fig. 4). Temperature significantly increased the float difference in salinity 25 ppt with a mean (±standard error (SE)) float difference of 0.6 (±0.31), 0.9 (±0.37), 0.5 (±0.21), and 1.5 (±0.64) at 3°C, 7°C, 12°C, and 16 °C, respectively (Table 1; Fig. 4). The float difference at salinity 35 ppt was significantly decreased by temperature with mean (±SE) float differences of -0.08 (±0.08), -0.08 (±0.08), -0.31 (±0.18), and -1.08 (±0.31) at 3, 7, 12, and 16 °C, respectively (Table 1; Fig. 4).

**Fig. 4.**
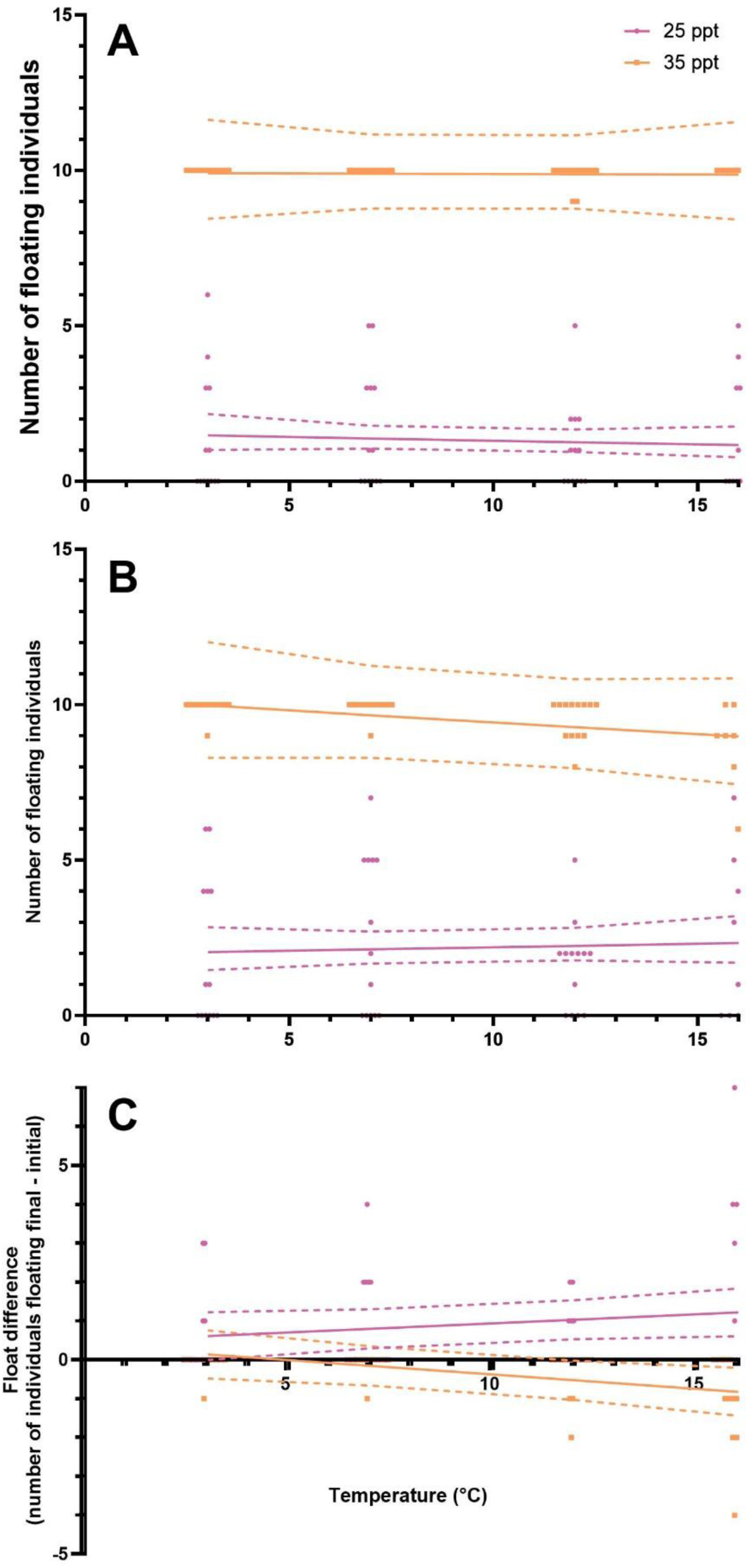
The number of individuals floating initially (A; Table d) and finally (B; Table c) was significantly different between salinities of 25 ppt (pink circles) and 35 ppt (orange squares), but these salinities were not significantly different in float difference (C; Table 1). Temperature (°C) had a significant positive and negative effect on float difference in salinites of 25 ppt and 35 ppt, respectively (Table 1), but no effect on the number of individuals floating initially (Table d) or finally (Table c). Plotted points are the raw data points. The solid line is the predicted relationship between temperature and the given response variable from glmmTMBs (Equation 1). The dashed lines display the 2.5% to 97.5% confidence interval for the line of best fit. n=13 beakers for T03-S25, T03-S35, T07-S25, T12-S25, T12-S35, T16-S25, and T16-S35 and n=12 beakers for T07S35.

**Table 1.** The results from three general linear mixed models (Equation 1), including the estimates, confidence intervals, p-values and adjusted p-values, testing the impact of salinity and temperature on the number of individuals floating initially, the number of individuals floating finally and the float difference. The σ^2^ and Marginal R^2^ values are included for each model. N=103.

|  | Number of individuals<br>floating initially |  |  | Number of individuals<br>floating finally |  |  | Float difference |  |  |
| --- | --- | --- | --- | --- | --- | --- | --- | --- | --- |
| Predictors | estimate | p-value | adjusted<br>p-value | estimate | p-value | adjusted<br>p-value | estimate | p-value | adjusted p-value |
| Intercept | 2.29 | <0.001 | <0.001 | 2.32 | <0.001 | <0.001 | 0.35 | 0.350 | 0.511 |
| Temperature | 0.00 | 0.969 | 0.969 | -0.01 | 0.373 | 0.511 | -0.07 | **0.014** | **0.028** |
| Salinity | -1.85 | <0.001 | <0.001 | -1.64 | <0.001 | <0.001 | 0.11 | 0.807 | 0.880 |
| Temperature:<br>Salinity | -0.02 | 0.493 | 0.592 | 0.02 | 0.383 | 0.511 | 0.12 | **0.005** | **0.011** |
| $\sigma^2$ | | 0.00 | | | 0.01 | | | 0.057 | |
| Marginal $R^2$ | | NULL | | | NULL | | | 0.466 | |

Bolded p-values indicate significance.

There were 4 (0.8%), 69 (13.3%), 25 (4.8%), 20 (3.9%), and 17 (3.2%) individuals among 4, 26, 15, 7, and 6 total beakers that underwent stage changes under early floated, sunk developed, developed floated, float developed, and late floated categories, respectively, in salinity 25 ppt treatments (Fig. 5). In S35 18 (3.5%), 114 (22.5%), and 1 (0.2%) individuals among 13, 35, and 1 total beakers underwent stage changes under early sank, developed, and sunk categories, respectively (Fig. 5). Temperature had a significant positive effect on the sunk developed category in salinity 35 ppt (Table S3; Fig. 5), with the number of individuals increasing from 2 at 3°C to 30 at 16°C, and the developed category in salinity 35 ppt (Table S4; Fig. 5), with 41 more individuals at 16°C than at 3°C. There was no significant effect of temperature on any other stage changes in salinity 25 ppt (Table S3) and salinity 35 ppt (Table S4). No stage changes were significantly different than zero.

**Fig. 5.**
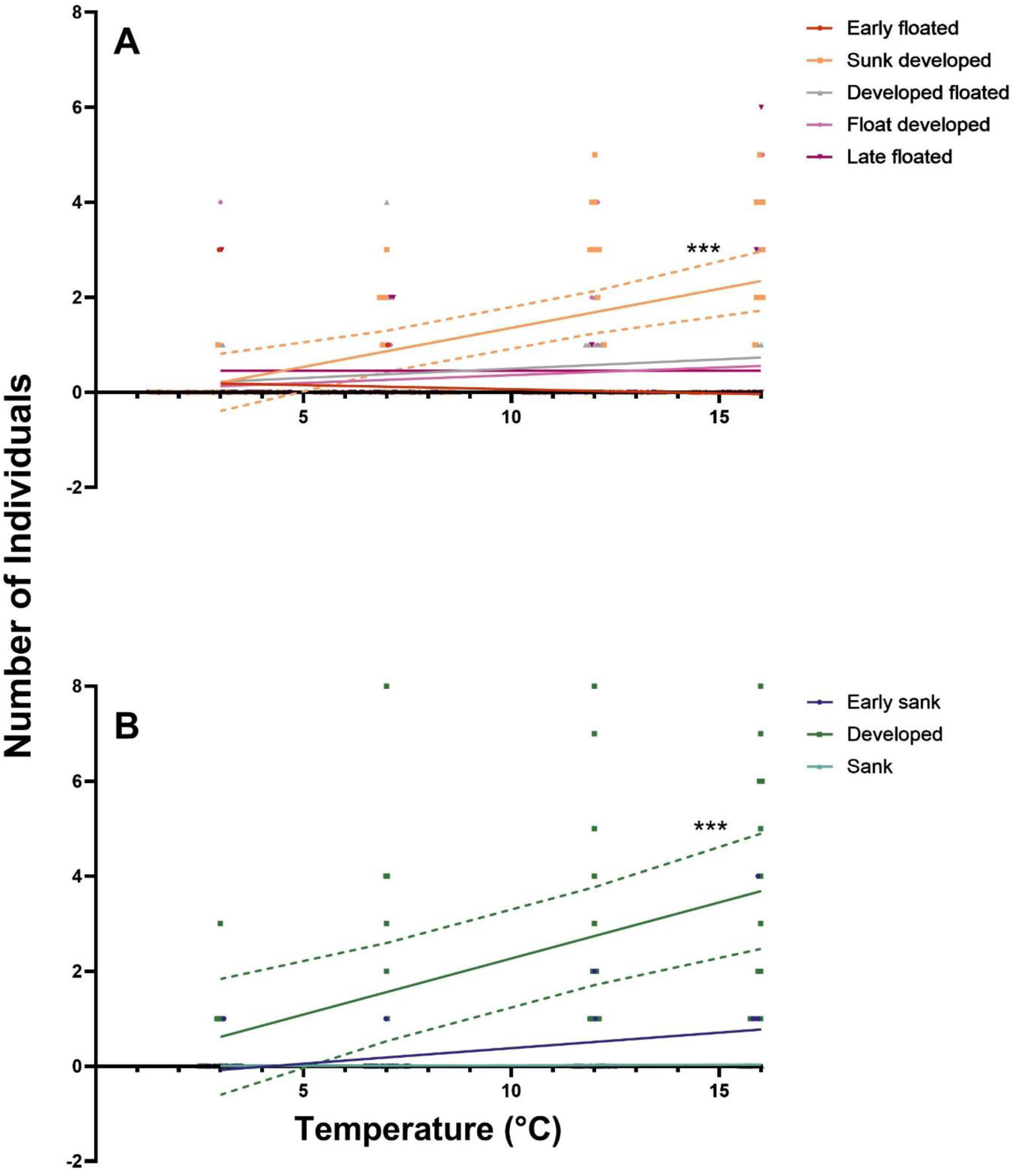
Temperature (°C) had a significant positive effect on the sunk developed (orange squares) and developed (green squares) stage changes in salinites of 25 ppt (A; Table S3) and 35 ppt (B; Table S4), respectively. The state changes that were not affected by temperature in 25 ppt salinity were early floated (red circles), developed floated (grey triangles), float developed (light purple diamonds) and late floated (dark purple inverted triangles). In 35 ppt salinity, stage changes of early sank (blue circles) and sank (teal triangles) were not affected by temperature. Plotted points are the raw data points. The solid line is the predicted relationship between temperature and the given stage change from glmmTMBs (Equation 2). The dotted lines display the 2.5% to 97.5% confidence interval for the line of best fit. Three stars indicate a significance less than 0.001. n=13 beakers for T03-S25, T03-S35, T07-S25, T12-S25, T12-S35, T16-S25, and T16-S35 and n=12 beakers for T07-S35.

### 3.3 The impact of inversions on survival

The first, second, and third dimensions of the PCA made with the differences in inversion genotype frequencies explained 38.74%, 29.52%, and 22.00% of the variation, respectively (Fig. S3). Inv12 genotypes were the largest explainer for dimension 1 (cos2=0.545), with the inv2 heterozygote (cos2=0.495) and derived (cos2=0.454) genotypes closely behind (Fig. S3). Inv7 and inv12 controlled most of dimensions 2 (cos2=0.566) and 3 (cos2=0.405) of the PCA axes, respectively (Fig. S3). Inv2 and inv12 grouped in a similar location in the PCA, indicating they are positively correlated, specifically, the heterozygotes and derived genotypes are positively correlated with each other (Fig. S3).

Neither salinity nor temperature impacted beakers’ locations along the first PCA axis (mean±SE is -0.246±0.399 and 0.436±0.292, for 25 and 35 ppt, respectively, and -1.19 (±0.489), 0.03 (±0.945) and 0.39 (±0.241) at 3°C, 7°C and 16°C, respectively; Table 2; Fig. 6). Temperature marginally significantly decreased beakers’ location along the third PCA axis, with the average (±SE) location being 0.088 (±0.346), 0.024 (±0.476), and -0.039 (±0.299) for 3°C, 7°C, and 16°C, respectively (Table 2; Fig. 6). Neither temperature nor salinity impacted a beaker’s location along the second PCA axis (Table 2; Fig. 6), with the average (±SE) location being 0.680 (±0.471), 0.338 (±0.445), -0.356 (±0.335), 0.321 (±0.310), and -0.569 (±0.350) for 3°C, 7°C, 16°C, 25 ppt, and 35 ppt, respectively. The beakers’ locations along the third PCA axis were significantly higher at 35 ppt (mean±SE is 0.585±0.304) compared to 25 ppt (mean ±SE is - 0.330 ±0.259; Table 2; Fig. 6).

**Fig. 6.**
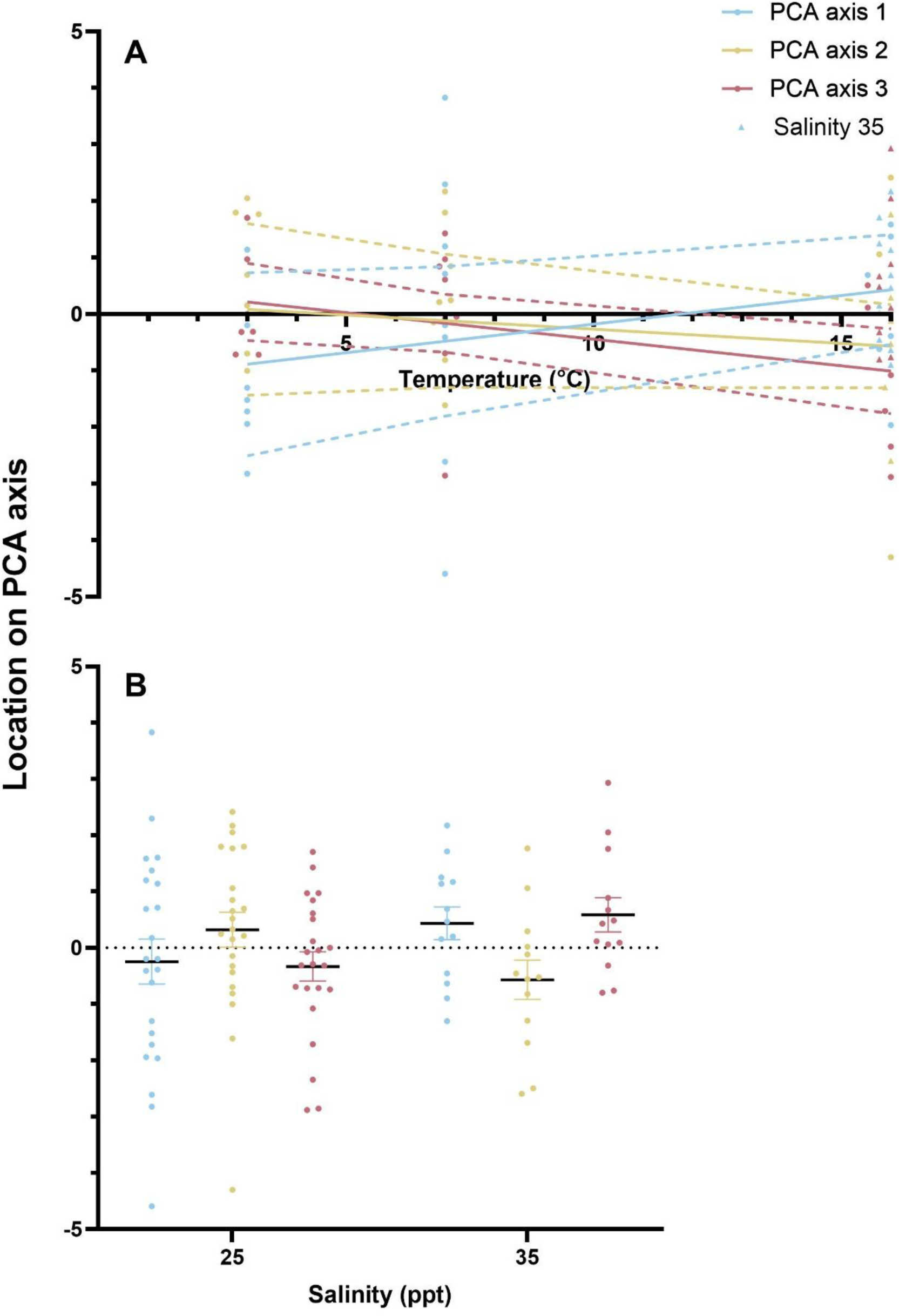
Temperature (°C; A) had a marginally significant positive and negative effect on the location of a beaker along PCA axis 1 (blue) and axis 3 (pink), respectively, while it did not affect axis 2 location (yellow). Salinity (ppt; B) significantly decreased the location of a beaker along PCA axis 3 and did not affect axes 1 and 2. Salinity 25 is plotted as circles, and salinity 35 is plotted as triangles in panel. **A.** Plotted points are the raw data points. The solid line is the predicted relationship between temperature and the given stage change from glmmTMBs (Equation 4). The dashed lines display the 2.5% to 97.5% confidence interval for the line of best fit. Error bars represent the standard error around the mean. n=7, 8, 8, and 13 beakers for T03-S25, T07-S25, T16-S25, and T16-S35, respectively.

**Table 2.** The results from three general linear mixed models (Equation 4), including the estimates, confidence intervals, p-values and adjusted p-values, testing the impact of salinity and temperature on the PCA axis 1, 2, and 3 locations. The PCA was created based on the genotype frequency differences. The σ^2^ and Marginal R^2^ values are included for each model. N=36.

|  | Axis 1 |  |  | Axis 2 |  |  | Axis 3 |  |  |
| --- | --- | --- | --- | --- | --- | --- | --- | --- | --- |
| Predictors | estimate | p-value | adjusted p-value | estimate | p-value | adjusted p-value | estimate | p-value | adjusted p-value |
| Intercept | -1.19 | 0.211 | 0.379 | 0.23 | 0.799 | 0.899 | 2.10 | **0.004** | **0.018** |
| Temperature | 0.10 | **0.046** | 0.102 | -0.05 | 0.336 | 0.477 | -0.09 | **0.022** | 0.066 |
| Salinity | 0.06 | 0.917 | 0.917 | 0.53 | 0.371 | 0.477 | -1.60 | **0.001** | **0.007** |
| $\sigma^2$ | 0.87 | | | 0.92 | | | 0.57 | | |
| Marginal $R^2$ | 0.258 | | | 0.205 | | | 0.397 | | |

Bolded p-values indicate significance.

Salinity had a significant negative impact on the ancestral allele frequency difference for inv12, with the average (±SE) decreasing from 0.0759 (±0.0123) at 25 ppt to 0.0170 (±0.0185) at 35 ppt (Table 3; Fig. 7). Temperature significantly impacted the ancestral allele frequency differences for inv2 (Table 3; mean±SE difference was 0.1315±0.0399, 0.0288±0.0779, and - 0.0395±0.0298 for 3°C, 7°C, and 16°C, respectively), but not the ancestral allele frequency differences for inv7 (Table 3; mean±SE difference was -0.0080 ±0.0266, -0.0246 ±0.0335, and - 0.0120 ±0.0118 for 3°C, 7°C, and 16°C, respectively), or inv12 (Table 3; mean ±SE difference was 0.0940±0.0187, 0.0567±0.0274, and 0.0407±0.0144 for 3°C, 7°C, and 16°C, respectively; Fig. 7). Salinity did not significantly impact the ancestral allele frequency differences for inv2 (Table 3; mean±SE difference was 0.0226±0.0407 and -0.0153±0.0237 for 25 and 25 ppt, respectively) and inv7 (Table 3; mean±SE difference was -0.0247±0.0152 and 0.0048±0.0133 for 25 and 25 ppt, respectively; Fig. 7).

**Fig. 7.**
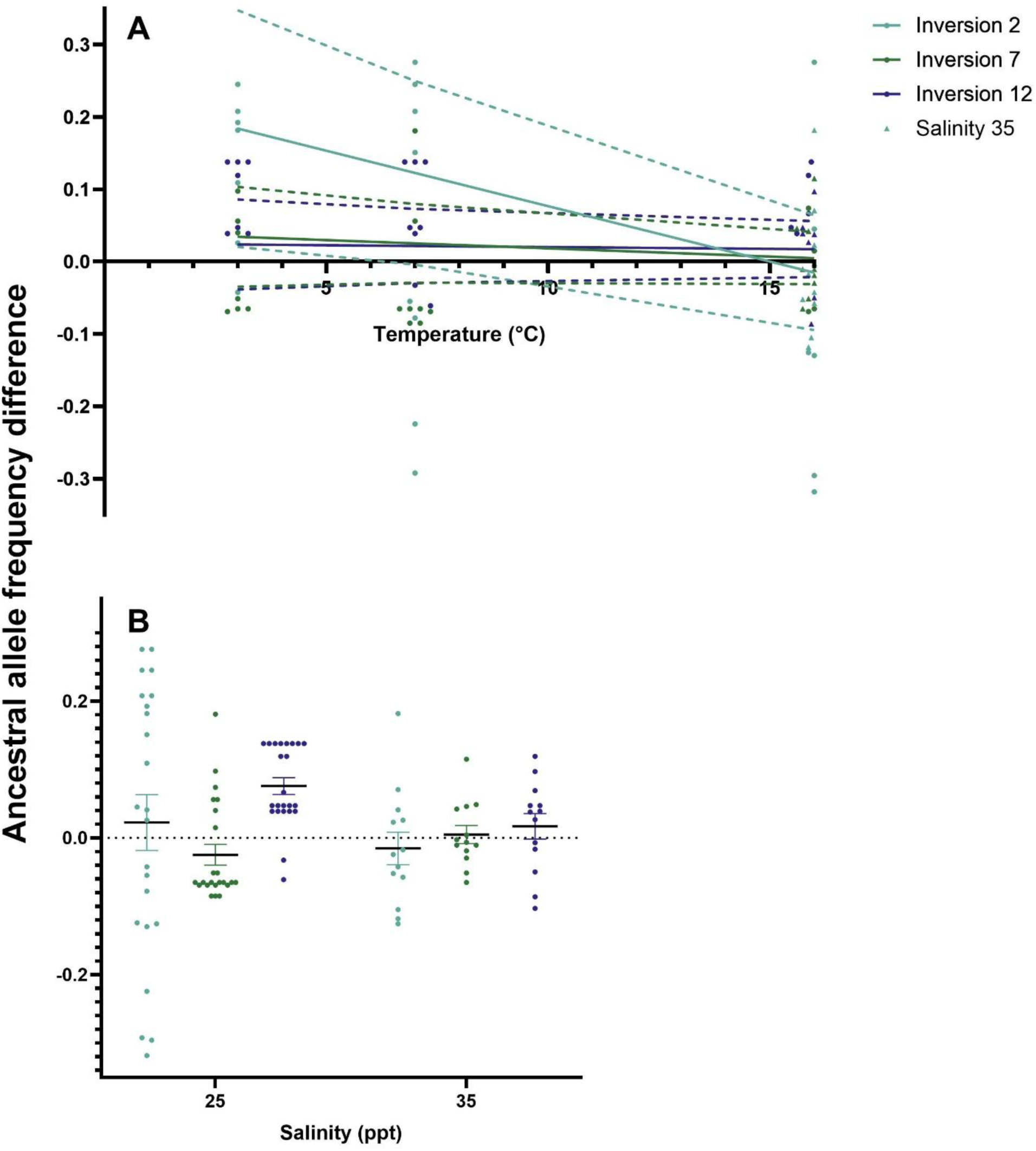
Temperature (°C; A) significantly impacted the ancestral allele frequency difference for inv2 (light blue), but not inv7 (green), or inv12 (dark blue). Salinity (ppt; B) significantly decreased the inv12ancestral allele frequency difference and did not affect inv2 or inv7 frequency differences. Plotted points are the raw data points. The solid line is the predicted relationship between temperature and the given stage change from glmmTMBs (Equation 3). Salinity 25 is plotted as circles, and salinity 35 is plotted as triangles in panel A. The dotted lines display the 2.5% to 97.5% confidence interval for the line of best fit. Error bars represent the standard error around the mean. n=7, 8, 8, and 13 beakers for T03-S25, T07-S25, T16-S25, and T16-S35, respectively.

**Table 3.** The results from three general linear mixed models (Equation 3), including the estimates, confidence intervals, p-values and adjusted p-values, testing the impact of salinity and temperature on the difference in the ancestral allele frequency for inv2, inv7 and inv12. The σ^2^ and Marginal R^2^ values are included for each model. N=36.

|  | Inversion 2 |  |  | Inversion 7 |  |  | Inversion 12 |  |  |
| --- | --- | --- | --- | --- | --- | --- | --- | --- | --- |
| Predictors | estimate | p-value | adjusted p-value | estimate | p-value | adjusted p-value | estimate | p-value | adjusted p-value |
| Intercept | 0.23 | **0.020** | 0.059 | 0.04 | 0.321 | 0.418 | 0.02 | 0.492 | 0.554 |
| Temperature | -0.02 | **0.006** | **0.041** | 0.00 | 0.326 | 0.418 | 0.00 | 0.798 | 0.798 |
| Salinity | -0.07 | 0.273 | 0.418 | -0.05 | 0.072 | 0.162 | 0.06 | **0.009** | **0.041** |
| $\sigma^2$ | | 0.01 | | | 0.00 | | | 0.00 | |
| Marginal $R^2$ | | 0.314 | | | 0.162 | | | 0.296 | |

Bolded p-values indicate significance.

### 3.4 Ecotype’s impact on survival

When comparing a batch’s probability of belonging to the offshore ecotype distribution to the distribution of the floating individuals within a given batch and treatment, there were no significant differences (Table S5).

## 4 Discussion

### 4.1 The effect of salinity and temperature on cod egg survival

Our results demonstrate that salinity has an immediate impact on egg buoyancy in Atlantic cod (*Gadus morhua*), influencing both initial and final floating status. As expected, a greater proportion of eggs sank at low salinity conditions (25 ppt) compared to ambient conditions (35 ppt). This reflects the disruption of neutral buoyancy, a critical factor for egg viability and survival in pelagic spawning species (Nissling and Westin, 1997; Schmidt et al., 2024). Our results are consistent with other studies which found that mortality (Schmidt et al., 2024) and hatching success (Bian et al., 2016; Schmidt et al., 2024) decrease in low salinity. When coupled with evidence that spermatogenesis is impaired under reduced salinity (Nissling and Westin, 1997), our results suggest that increasing freshwater input, such as that driven by changes in precipitation and runoff, could have negative consequences for cod reproductive success and population structure.

While temperature did not significantly affect the initial or final floating state of eggs, it did influence the difference in buoyancy over the 24-hour period. Specifically, floatation stability declined as temperature increased, with the float difference diverging further from zero at higher temperatures. We did not observe a detrimental effect at low temperature (3°C) on egg survival over the short exposure period, contrasting with previous studies reporting reduced survival under similar low-temperature conditions through development (Jordaan and Kling, 2003). These studies used populations that spawn at different times, and survival in stressful temperatures corresponds with spawning time (Oomen and Hutchings, 2015; Pepin, 1991; Geffen et al., 2006). Our finding that survival significantly decreased at higher temperatures above 10°C is consistent with the literature (Hasan et al., 2023; Geffen et al., 2006), indicating that continued increases in ocean temperatures are likely to majorly impact cod egg survival.

Salinity interacted with temperature to impact float difference particularly at high temperatures. Salinity controlled the direction in which the float difference occurred. In low salinity, there is a positive float difference, and at high salinity, there is a negative float difference. Similarly, in both Pacific cod and Atlantic cod, long-term salinity stress exposure only impacted egg phenotype at the edges of thermal tolerance (Bian et al., 2016). However, a study that applied both temperature and salinity stress on eggs over the entirety of development did not find any interaction between salinity and temperature (Laurence and Rogers, 1976). The difference between our results and this study might be due to the adaptation of both temperature and salinity being seasonally dependent (Perry et al., 2024; Magill and Sayer, 2004; Pepin, 1991). Overall, temperature is the main stressor causing mortality in eggs under both acute and long-term conditions (Bian et al., 2016).

Interestingly, some individuals who initially sank at 25 ppt floated after 24 hours (hereafter, termed ‘recovery’). Recovery likely occurred because of osmosis through the oocyte plasma membrane, which decreased the egg salinity at neutral buoyancy, allowing it to float at 25 ppt (Fridman, 2020). Osmosis through the oocyte plasma membrane occurs quickly. Individuals who recovered likely had a neutral buoyancy just above 25 ppt initially because osmosis would only minimally adjust egg buoyancy. The same trend, of low salinity water decreasing neutral buoyancy, is seen in cod yolk-sac larvae (Schmidt et al., 2024). More recovery was also seen at higher temperatures, which might indicate that recovery is faster at higher temperatures or that buoyancy changes are linked to life stage changes correlated with temperature.

### 4.2 The effect of temperature on life stage changes

More individuals developed at higher temperatures, which is common in fishes (Pepin, 1991; Slesinger et al., 2024), including Atlantic cod (Jordaan and Kling, 2003; Otterlei et al., 1999; Righton et al., 2010). The temperature at which cod growth rate decreases varies among populations (Righton et al., 2010; Oomen and Hutchings, 2015, 2016; Oomen et al., 2022). We observed an increase in development between 12°C and 16°C on an acute time scale. We did not find a significant effect of temperature on any other life stage categories.

### 4.3 The effect of salinity and temperature on inversion frequencies

An increase in temperature decreased the frequency of the inv2 ancestral allele and selected for the inv2 and inv12 derived genotypes. Derived inv2 and inv12 genotypes might contribute to warm temperature adaptation. Our results align with multiple studies that correlated these inversions with temperature (Barney et al., 2017; Puncher et al., 2019; Berg et al., 2015; Johansen et al., 2020), specifically indicating that these inversions facilitate cold adaptation (Oomen, 2019). Due to there being minimal mortality in cold temperatures and few or no inv2 and inv12 ancestral individuals, it is unclear whether the ancestral genotype is cold-adapted. However, we did not see selection against the derived genotype or the derived allele frequency at cold temperatures. Therefore, ancestral genotypes and ancestral allele frequencies increasing in low temperatures are likely due to selection for the ancestral genotype and the ancestral allele and not against the derived genotypes or allele B. There was no effect of temperature on the ancestral allele frequency for inv12. This might indicate that temperature affects inv12 genotypes and not alleles, or that the temperature effect is only present due to correlations between inv2 and inv12 genotypes.

Contrary to previous research (Berg et al., 2015; Barth et al., 2017), we did not observe an effect of salinity on the inv2 ancestral allele frequency or inv2 genotypes. It is plausible that inv2 impacts salinity adaptation in the larval stage but not the egg stage. During the egg stage, fish cannot control water and ion movement between the egg sac and the external environment (Fridman, 2020). After hatching, ionocytes help larvae osmoregulate until the gills are fully formed (Fridman, 2020). Therefore, genes contained within inv2 might control osmoregulation in larvae and later life stages, helping these life stages, but not the egg stage, to adapt to low-salinity conditions (Berg et al., 2015).

Inv12 derived allele and derived genotypes (based on PC3) were selected against in low salinity. Salinity adaptation is correlated with inv2 more often than inv12 (Barth et al., 2017; Berg et al., 2015; Oomen, 2019), with only one correlative (Sodeland et al., 2016) and one gene annotation (Sodeland et al, 2022) study linking inv12 and low salinity. However, we show that inv12 is linked with low salinity at the egg stage. Genes contained in inv12 might contribute to buoyancy.

The allele and genotype (PC2) frequencies of inv7 were not affected by temperature or salinity. This result was not expected, as previous studies found genes controlling low salinity adaptation in inv7 (Sodeland et al, 2022). Inv7 genotypes are also correlated with temperature, both in the wild (Johansen et al., 2020) and in laboratory experiments (Oomen, 2019). It is possible that genes within inv7 are not activated at the egg stage or under acute time frames. There were no inv7-derived individuals in our data set.

Inversion genotype frequencies revealed that inv2 and inv12 were correlated with each other. Specifically, heterozygote genotypes and derived genotypes were correlated between inv2 and inv12. Both inv2 and inv12 genotype frequencies were impacted by temperature, which might indicate that inv2 and inv12 are interacting with each other in temperature adaptation. Inv7 controlled a separate axis on the PCA and was uncorrelated with inv2 or inv12. Multiple studies found that more than one inversion correlated with both salinity and temperature when researching them at the same time (Sodeland et al., 2016; 7; Sodeland et al, 2022; Johansen et al., 2020). Experimental studies that breed for specific inversion genotypes are needed to disentangle inversion correlations.

### 4.4 The role of random effects and ecotype in salinity and temperature adaptation

There was high variance (above 0.50) of the beaker nested in the batch random effect in the models that analyzed the float difference and PCA axes locations. The high variance was mostly due to differences between batches, consistent with a previous study that found timing of sampling impacted cod egg phenotype (Jordaan et al., 2006). For example, in batches 3 and 4, few individuals floated throughout compared to the other three batches (Fig. S4). One possible reason for the difference between the batches was the collection date conditions. For example, rain occurred prior to batch 3 collection. The water in the top 0.5 meters of the pond was fresher than the ambient conditions. Individuals with lower neutral buoyancy were exposed to fresh surface conditions and were not frequent in batch 3, potentially due to high mortality in the pond. This mortality might have been caused by high water movement. Similarly, there were high winds during the collection of batch 4. We saw that more eggs in batch 4b floated, indicating that individuals with higher neutral buoyancies were blown into the corner of the pond.

### 4.5 Conclusion

We used common garden experiments to dissentangle the contributions of chromosomal inversions from other genomic regions in shaping stressor tolerance. We provide experimental evidence suggesting a role of inv2 in temperature adaptation and inv12 in salinity adaptation in Atlantic cod eggs. The correlations between inversions should be further researched. Low salinity and high temperatures are stressful for egg-staged cod even on a short time frame, indicating that there is potential for extreme die-off events during low salinity input and high temperature events. These results might explain why Atlantic cod is having trouble recovering, but fishing pressure still needs to be reduced (Hilborn and Litzinger, 2009). To prevent future high temperatures and freshening stress impacting cod populations, further global warming needs to be prevented by reducing fossil fuel output (IPCC, 2023).

## Acknowledgements

We respectfully acknowledge the research conducted at the University of New Brunswick, SJ was conducted on unsurrendered and unceded traditional Wəlastəkwiyik (Wolastoqiyik) and Mi’kmaq (part of the Wabanaki) land. This territory is part of the Peace and Friendship Treaties, which did not involve the surrender of lands, waters, or resources and established an ongoing relationship of peace, friendship and mutual respect between equal nations. Tusen takk til Stian Stiansen for hjelpen med eksperimentene. Individuals at the Tjärnö Marine Laboratory and Flødevigen Research Station, Institute of Marine Research, particularly Hanne Sannæs and Heidi Fiskaa, are thanked for all their help. Thanks to Hannah Earp and the GEcoKelp project for their loan of the experimental aquarium heat blocks that were used in this study. Thanks go out to Dr. Scott Pavey and Dr. Alex Zimmer at UNBSJ for their guidance on the project. We’d like to thank Claudia Lacroix, Abigail K. Scher, and Stephanie Maheux for the feedback that they provided.

## Data and resource availability

Data is deposited in the Mendeley Data repository doi: https://10.17632/46xbmvcg6h.1

## Appendix A: Pilot Experiments

### 1.0 General Methods

All methodology and equipment not explicitly stated followed methodology in the Materials and Methods section. Pilot experiments took place during the week of February 3rd, 2025.

### 2.0 Temperature trial

#### 2.1 Methods

We placed individuals in beakers with 200 ml of water at four different temperatures: <1°C (n=7), 3°C (n=10), 14°C (n=6), and 16°C (n=6). We monitored beakers for up to 48 hours to help determine which temperatures caused sinking and when. Beakers were mixed each time sunk/float measurements were taken. For temperatures 3°C, 14°C, and 16°C, we used the temperature gradient block with the 3°C, 14°C, and 16°C beakers located in columns K, C, and A, respectively. To create the <1°C treatment, we used an ice bath with freshwater ice frozen at - 20°C and no water. We moved the ice in the container closer to the beaker when needed, but there was no other manipulation of the ice bath. Temperature in the <1°C trial increased steadily throughout the 24 hours that individuals were in the treatment, but did not increase past 3°C.

#### 2.2 Results

In the <1°C and 3°C treatments, no individuals sank throughout the experiments. In the first 8 hours of treatment, no individuals sank in the 14°C and 16°C treatments; some individuals sank after 19 hours, with the majority of individuals sinking in the first 24 hours (Fig. S5).

#### 2.3 Discussion

Due to there being no difference in the <1°C and 3°C treatments, 3°C was chosen as the cold temperature because we were able to stabilize 3°C for 24 hours in the temperature gradient block. We chose to exclude the 14°C treatment and include the 12°C treatment because 12°C is seen as the upper end of the temperature at which cod spawn (Righton et al., 2010). Therefore, results on the ability of cod eggs to survive stress at 12°C were more biologically interesting than a 14°C treatment. We chose to apply treatments for 24 hours because there was minimal sinking that occurred after 24 hours, meaning the first 24 hours of exposure were crucial for survival.

### 3.0 Salinity trial

#### 3.1 Methods

We ran salinity trials with four treatments: 10, 20, 25, and 30 ppt. Individuals were placed in a 250 ml beaker with 200 ml of low salinity water at the respective salinity. We only observed the immediate response to the 10 and 20 ppt treatments. The 25 and 30 ppt trials had replicates at two temperatures, 3°C (n=8) and 16°C (n=7) for 25 ppt and 7°C (n=11) and 16°C (n=9) for 30 ppt, with the temperature being controlled by the thermal gradient block. We monitored the number of individuals that sank and floated for 24 and 48 hours in the 25 and 30 ppt trials, respectively. We stirred the beakers each time a measurement was taken.

#### 3.2 Results

All individuals in the 10 and 20 ppt treatments sank immediately. After 24 hours in 25 ppt at 3°C, there was no change in the number of individuals that sank, while at 16°C, 1 individual sank after 3 hours. In 30 ppt at 7°C, 3 individuals sank between 23 and 28 hours. Finally, in the 30 ppt treatment at 16°C, 3 individuals sank after 20 hours, and 1 additional individual sank after 44 hours (Fig. S6).

#### 3.3 Discussion

We chose to use 25 ppt as the low salinity treatment for the temperature and salinity stressor experiment because it was the salinity at which individuals both sank and floated immediately. At 20 ppt and lower, all individuals sank immediately, and at 30 ppt, all individuals floated immediately. Therefore, at 25 ppt, there is the potential for individuals to change their state.

### 4.0 Dessication trial

#### 4.1 Methods

We completed desiccation trials to determine how long individuals could survive in the beakers without being stirred. Three treatments were applied: 22, 24 and 48 hours of dessication. Beakers with normal salinity water were placed in the thermal gradient block stabilized at 7°C. 6, 8, and 10 individuals were placed in the 22, 24, and 48-hour treatments, respectively. After the determined amount of time, we stirred the beakers to move individuals stuck to the surface or the side of the beaker. Then, we counted the number of individuals sunk and floating.

#### 4.2 Results

After 22, 24, and 48 hours without stirring, 0 out of 6, 1 out of 8, and 2 out of 10 individuals sank, respectively.

#### 4.3 Discussion

Due to there being minimal sinking after 24 hours of dessication and no observed sinking after 22 hours of dessication, it was determined that stirring individuals once during treatments, which only exposed them to 12 hours of dessication, would not have negative impacts on survival in the temperature and salinity stressor experiment. This indicates that cod eggs have a good ability to withstand dessication over short periods and that dessication is not a concern for short-term stress experiments.

## References

Ackerly, K. L., Roark, K. J. and Nielsen, K. M. (2023). Short-term salinity stress during early development impacts the growth and survival of red drum (*Sciaenops ocellatus*). Estuaries Coast. 46, 541–550.

Andersson, L., Bekkevold, D., Berg, F., Farrell, E. D., Felkel, S., Ferreira, M. S., Fuentes-Pardo, A. P., Goodall, J. and Pettersson, M. (2024). How fish population genomics can promote sustainable fisheries: A road map. Annu. Rev. Anim. Biosci. 12, 1–20.

Árnason, T., Magnadóttir, B., Björnsson, B., Steinarsson, A. and Björnsson, B. T. (2013). Effects of salinity and temperature on growth, plasma ions, cortisol and immune parameters of juvenile Atlantic cod (*Gadus morhua*). Aquaculture 380–383, 70–79.

Barney, B. T., Munkholm, C., Walt, D. R. and Palumbi, S. R. (2017). Highly localized divergence within supergenes in Atlantic cod (*Gadus morhua*) within the Gulf of Maine. BMC Genomics 18, 271.

Barrett, G. W., van Dyne, G. M. and Odum, E. P. (1976). Stress Ecology. Bioscience 26, 192– 194.

Barth, J. M. I., Berg, P. R., Jonsson, P. R., Bonanomi, S., Corell, H., Hemmer-Hansen, J., Jakobsen, K. S., Johannesson, K., Jorde, P. E., Knutsen, H., et al. (2017). Genome architecture enables local adaptation of Atlantic cod despite high connectivity. Mol. Ecol. 26, 4452–4466.

Berg, P. R., Jentoft, S., Star, B., Ring, K. H., Knutsen, H., Lien, S., Jakobsen, K. S. and André, C. (2015). Adaptation to low salinity promotes genomic divergence in Atlantic cod (*Gadus morhua* L.). Genome Biol. Evol. 7, 1644–1663.

Bian, X., Zhang, X., Sakurai, Y., Jin, X., Wan, R., Gao, T. and Yamamoto, J. (2016). Interactive effects of incubation temperature and salinity on the early life stages of pacific cod *Gadus macrocephalus*. Deep Sea Res. Part 2 Top. Stud. Oceanogr. 124, 117–128.

Bradbury, I. R., Hubert, S., Higgins, B., Borza, T., Bowman, S., Paterson, I. G., Snelgrove, P. V. R., Morris, C. J., Gregory, R. S., Hardie, D. C., et al. (2010). Parallel adaptive evolution of Atlantic cod on both sides of the Atlantic Ocean in response to temperature. Proc. Biol. Sci. 277, 3725–3734.

Brooks, M., Kristensen, K., Benthem, K. van, Magnusson, A., Berg, C., Nielsen, A., Skaug, H., Mächler, M. and Bolker, B. (2017). GlmmTMB balances speed and flexibility among packages for zero-inflated generalized linear mixed modeling. R J. 9, 378.

Calvin, K., Dasgupta, D., Krinner, G., Mukherji, A., Thorne, P. W., Trisos, C., Romero, J., Aldunce, P., Barret, K., Blanco, G., et al. (2023). IPCC, 2023: Climate Change 2023: Synthesis Report, summary for Policymakers. Contribution of working groups I, II and III to the Sixth Assessment Report of the Intergovernmental Panel on Climate Change [core writing team, H. lee and J. romero (eds.)]. IPCC, Geneva, Switzerland. *IPCC, 2023: Climate Change 2023: Synthesis Report. Contribution of Working Groups I, II and III to the Sixth Assessment Report of the Intergovernmental Panel on Climate Change [Core Writing Team, H. Lee and J. Romero (eds.)]. IPCC, Geneva, Switzerland* 1–34.

Catalán, I. A., Reglero, P. and Álvarez, I. (2020). Research on early life stages of fish: a lively field. Mar. Ecol. Prog. Ser. 650, 1–5.

Cayuela, H., Rougemont, Q., Laporte, M., Mérot, C., Normandeau, E., Dorant, Y., Tørresen, O. K., Hoff, S. N. K., Jentoft, S., Sirois, P., et al. (2019). Standing genetic variation and chromosomal rearrangements facilitate local adaptation in a marine fish. bioRxiv 782201.

Chen K, Marschall EA, Sovic MG, Fries AC, Gibbs HL, Ludsin SA (2024). assignPOP: Population Assignment using Genetic, Non-Genetic or Integrated Data in a Machine Learning Framework. R package version 1.3.0.

Conover, W. J. (1972). A kolmogorov goodness-of-fit test for discontinuous distributions. J. Am. Stat. Assoc. 67, 591.

Côté, I. M., Darling, E. S. and Brown, C. J. (2016). Interactions among ecosystem stressors and their importance in conservation. Proc. Biol. Sci. 283, 20152592.

Cyr, F. and Galbraith, P. S. (2021). A climate index for the Newfoundland and Labrador shelf. *Earth Syst*. Sci. Data 13, 1807–1828.

DFO (2023). Stock Status Update of Atlantic Cod (Gadus morhua) in NAFO Divisions 4X5Y for 2022. DFO Can. Sci. Advis. Sec. Sci. Resp. Resp. 2023/017,.

Dray, S. and Dufour, A.-B. (2007). Theade4Package: Implementing the duality diagram for ecologists. J. Stat. Softw. 22.

Dutil, J.-D., Munro, J., Audet, C. and Besner, M. (1992). Seasonal variation in the physiological response of Atlantic cod (*Gadus morhua*) to low salinity. Can. J. Fish. Aquat. Sci. 49, 1149–1156.

Fang, Z., Pyhäjärvi, T., Weber, A. L., Dawe, R. K., Glaubitz, J. C., González, J. de J. S., Ross-Ibarra, C., Doebley, J., Morrell, P. L. and Ross-Ibarra, J. (2012). Megabase-scale inversion polymorphism in the wild ancestor of maize. Genetics 191, 883–894.

Fransson, A., Chierici, M., Nomura, D., Granskog, M. A., Kristiansen, S., Martma, T. and Nehrke, G. (2015). Effect of glacial drainage water on the CO_2_ system and ocean acidification state in an Arctic tidewater-glacier fjord during two contrasting years. J. Geophys. Res. Oceans 120, 2413–2429.

Fridman, S. (2020). Ontogeny of the osmoregulatory capacity of teleosts and the role of ionocytes. Front. Mar. Sci. 7.

Geffen, A. J., Fox, C. J. and Nash, R. D. M. (2006). Temperature-dependent development rates of cod *Gadus morhua* eggs. J. Fish Biol. 69, 1060–1080.

Gruber, B., Unmack, P. J., Berry, O. F. and Georges, A. (2018). dartr: Anrpackage to facilitate analysis of SNP data generated from reduced representation genome sequencing. Mol. Ecol. Resour. 18, 691–699.

Hamilton, L. C., Haedrich, R. L. and Duncan, C. M. (2004). Above and below the water: Social/ecological transformation in northwest Newfoundland. Popul. Environ. 25, 195–215.

Hemmer-Hansen, J., Nielsen, E. E., Therkildsen, N. O., Taylor, M. I., Ogden, R., Geffen, A. J., Bekkevold, D., Helyar, S., Pampoulie, C., Johansen, T., et al. (2013). A genomic island linked to ecotype divergence in Atlantic cod. Mol. Ecol. 22, 2653–2667.

Hilborn, R. and Litzinger, E. (2009). Causes of decline and potential for recovery of Atlantic cod populations. Open Fish Sci. J. 2, 32–38.

Hoffmann, A. A., Sgrò, C. M. and Weeks, A. R. (2004). Chromosomal inversion polymorphisms and adaptation. Trends Ecol. Evol. 19, 482–488.

Hollenbeck, C. M., Portnoy, D. S., Garcia de la Serrana, D., Magnesen, T., Matejusova, I. and Johnston, I. A. (2022). Temperature-associated selection linked to putative chromosomal inversions in king scallop (Pecten maximus). Proc. Biol. Sci. 289, 20221573.

IUCN (1996). Gadus morhua: Sobel, J. IUCN Red List of Threatened Species.

Johansen, T., Besnier, F., Quintela, M., Jorde, P. E., Glover, K. A., Westgaard, J.-I., Dahle, G., Lien, S. and Kent, M. P. (2020). Genomic analysis reveals neutral and adaptive patterns that challenge the current management regime for East Atlantic cod *Gadus morhua* L. Evol. Appl. 13, 2673–2688.

Jordaan, A. and Kling, L. J. (2003). Determining the optimal temperature range for Atlantic cod (*Gadus morhua*) during early life. The Big Fish Bang. In Proceedings of the 26th Annual Larval Fish Conference.

Jordaan, A., Hayhurst, S. E. and Kling, L. J. (2006). The influence of temperature on the stage at hatch of laboratory reared *Gadus morhua* and implications for comparisons of length and morphology. J. Fish Biol. 68, 7–24.

Kirkpatrick, M. (2010). How and why chromosome inversions evolve. PLoS Biol. 8, e1000501.

Kjesbu, O. S., Witthames, P. R., Solemdal, P. and Greer Walker, M. (1998). Temporal variations in the fecundity of Arcto-Norwegian cod (*Gadus morhua*) in response to natural changes in food and temperature. J. Sea Res. 40, 303–321.

Lambert, Y., Dutil, J.-D. and Munro, J. (1994). Effects of intermediate and low salinity conditions on growth rate and food conversion of Atlantic cod (*Gadus morhua*). Can. J. Fish. Aquat. Sci. 51, 1569–1576.

Laurence, G. C. and Rogers, C. A. (1976). Effects of temperature and salinity on comparative embryo development and mortality of Atlantic cod (*Gadus morhua* L.) and haddock (*Melanogrammus aeglefinus* (L.)). ICES J. Mar. Sci. 36, 220–228.

Link, J. S., Bogstad, B., Sparholt, H. and Lilly, G. R. (2009). Trophic role of Atlantic cod in the ecosystem. Fish Fish. 10, 58–87.

Lüdecke, D. (2018). Ggeffects: Tidy data frames of marginal effects from regression models. J. Open Source Softw. 3, 772.

Magill, S. H. and Sayer, M. D. J. (2004). The effect of reduced temperature and salinity on the blood physiology of juvenile Atlantic cod. J. Fish Biol. 64, 1193–1205.

Mérot, C., Oomen, R. A., Tigano, A. and Wellenreuther, M. (2020). A roadmap for understanding the evolutionary significance of structural genomic variation. Trends Ecol. Evol. 35, 561–572.

Mérot, C., Berdan, E. L., Cayuela, H., Djambazian, H., Ferchaud, A.-L., Laporte, M., Normandeau, E., Ragoussis, J., Wellenreuther, M. and Bernatchez, L. (2021). Locally adaptive inversions modulate genetic variation at different geographic scales in a seaweed fly. Mol. Biol. Evol. 38, 3953–3971.

Mohibul Hasan, M., Sultana Mely, S., Faruk, A. and Nayeem Hossain, M. (2023). Climate change effects on hatching success, embryonic development and larvae survival of freshwater fish: A critical review. Development 16, 24.

Nagelkerken, I., Allan, B. J. M., Booth, D. J., Donelson, J. M., Edgar, G. J., Ravasi, T., Rummer, J. L., Vergés, A. and Mellin, C. (2023). The effects of climate change on the ecology of fishes. PLOS Clim. 2, e0000258.

Nielsen, E. E., Hemmer-Hansen, J., Larsen, P. F. and Bekkevold, D. (2009). Population genomics of marine fishes: identifying adaptive variation in space and time. Mol. Ecol. 18, 3128–3150.

Nissling, A. and Westin, L. (1997). Salinity requirements for successful spawning of Baltic and Belt Sea cod and the potential for cod stock interactions in the Baltic Sea. Mar. Ecol. Prog. Ser. 152, 261–271.

Oomen, R. A. (2019). THE GENOMIC BASIS AND SPATIAL SCALE OF VARIATION IN THERMAL RESPONSES OF ATLANTIC COD (GADUS MORHUA).

Oomen, R. A. and Hutchings, J. A. (2015). Variation in spawning time promotes genetic variability in population responses to environmental change in a marine fish. Conserv. Physiol. 3, cov027.

Oomen, R. A., and Hutchings, J. A. (2016). Genetic variation in plasticity of life-history traits between Atlantic cod (*Gadus morhua*) populations exposed to contrasting thermal regimes. Can. J. Zoo. 94(4), 257–264.

Oomen, R. A., Kuparinen, A. and Hutchings, J. A. (2020). Consequences of single-locus and tightly linked genomic architectures for evolutionary responses to environmental change. J. Hered. 111, 319–332.

Orr, J. A., Macaulay, S. J., Mordente, A., Burgess, B., Albini, D., Hunn, J. G., Restrepo-Sulez, K., Wilson, R., Schechner, A., Robertson, A. M., et al. (2024). Studying interactions among anthropogenic stressors in freshwater ecosystems: A systematic review of 2396 multiple-stressor experiments. Ecol. Lett. 27, e14463.

Orr, J. A., Vinebrooke, R. D., Jackson, M. C., Kroeker, K. J., Kordas, R. L., Mantyka-Pringle, C., Van den Brink, P. J., De Laender, F., Stoks, R., Holmstrup, M. and Matthaei, C. D. (2020). Towards a unified study of multiple stressors: divisions and common goals across research disciplines. Proc. R. Soc. B. 287, 20200421.

Otterlei, E., Nyhammer, G., Folkvord, A. and Stefansson, S. O. (1999). Temperature- and size-dependent growth of larval and early juvenile Atlantic cod (*Gadus morhua*): a comparative study of Norwegian coastal cod and northeast Arctic cod. Can. J. Fish. Aquat. Sci. 56, 2099– 2111.

Pepin, P. (1991). Effect of temperature and size on development, mortality, and survival rates of the pelagic early life history stages of marine fish. Can. J. Fish. Aquat. Sci. 48, 503–518.

Pepin, P., Orr, D. C. and Anderson, J. T. (1997). Time to hatch and larval size in relation to temperature and egg size in Atlantic cod (*Gadus morhua*). Can. J. Fish. Aquat. Sci. 54, 2–10.

Perry, D., Tamarit, E., Sundell, E., Axelsson, M., Bergman, S., Gräns, A., Gullström, M., Sturve, J. and Wennhage, H. (2024). Physiological responses of Atlantic cod to climate change indicate that coastal ecotypes may be better adapted to tolerate ocean stressors. Sci. Rep. 14, 12896.

Puncher, G. N., Rowe, S., Rose, G. A., Leblanc, N. M., Parent, G. J., Wang, Y., and Pavey, S. A. (2019). Chromosomal inversions in the Atlantic cod genome: Implications for management of Canada’s Northern cod stock. Fish. Res. 216, 29–40.

R Core Team. (2023). R: A Language and Environment for Statistical Computing. R Foundation for Statistical Computing, *Vienna, Austria*. <https://www.R-project.org/>.

Righton, D. A., Andersen, K. H., Neat, F., Thorsteinsson, V., Steingrund, P., Svedäng, H., Michalsen, K., Hinrichsen, H. H., Bendall, V., Neuenfeldt, S., et al. (2010). Thermal niche of Atlantic cod *Gadus morhua*: limits, tolerance and optima. Mar. Ecol. Prog. Ser. 420, 1–13.

Schmidt, N., Garate-Olaizola, M. and Laurila, A. (2024). Acclimatizing laboratory-reared hatchling cod (*Gadus morhua*) to salinity conditions in the Baltic Sea. Aquaculture 579, 740255.

Shapero, M. (2013). SNP Genotyping using Affymetrix’ Axiom® Genotyping Solution.

Sinclair-Waters, M., Bradbury, I. R., Morris, C. J., Lien, S., Kent, M. P. and Bentzen, P. (2018). Ancient chromosomal rearrangement associated with local adaptation of a postglacially colonized population of Atlantic Cod in the northwest Atlantic. Mol. Ecol. 27, 339–351.

Slesinger, E., Mundorff, S., Laurel, B. J. and Hurst, T. P. (2024). The combined effects of ocean warming and ocean acidification on Pacific cod (*Gadus macrocephalus*) early life stages. Mar. Biol. 171.

Sodeland, M., Jorde, P. E., Lien, S., Jentoft, S., Berg, P. R., Grove, H., Kent, M. P., Arnyasi, M., Olsen, E. M. and Knutsen, H. (2016). “Islands of divergence” in the Atlantic cod genome represent polymorphic chromosomal rearrangements. Genome Biol. Evol. 8, 1012–1022.

Sodeland, M., Jentoft, S., Jorde, P. E., Mattingsdal, M., Albretsen, J., Kleiven, A. R., Synnes, A.-E. W., Espeland, S. H., Olsen, E. M., Andrè, C., et al. (2022). Stabilizing selection on Atlantic cod supergenes through a millennium of extensive exploitation. Proc. Natl. Acad. Sci. U. S. A. 119, e2114904119.

Tesdal, J.-E. and Haine, T. W. N. (2020). Dominant terms in the freshwater and heat budgets of the subpolar north Atlantic ocean and Nordic seas from 1992 to 2015. J. Geophys. Res. Oceans 125.

Thompson, B. M. and Riley, J. D. (1981). Egg and larval development studies in the North Sea cod (*Gadus morhua* L.). Rapp. P.-v. Réun. Cons. Int. Explor. Mer. 178, 553–559.

Veenhof, R. J., Champion, C., Dworjanyn, S. A., Schwoerbel, J., Visch, W. and Coleman, M. A. (2024). Projecting kelp (*Ecklonia radiata*) gametophyte thermal adaptation and persistence under climate change. Ann. Bot. 133, 153–168.

Vega-Trejo, R., Head, M. L., Jennions, M. D. and Kruuk, L. E. B. (2018). Maternal-by-environment but not genotype-by-environment interactions in a fish without parental care. Heredity 120, 154–167.

Wellenreuther, M. and Bernatchez, L. (2018). Eco-evolutionary genomics of chromosomal inversions. Trends Ecol. Evol. 33, 427–440.

Wellenreuther, M., Oomen, R. A., Young Han, K., Krohman, R., and T. B. H. Reusch. (2025). Beyond supergenes: the diverse roles of inversions in trait evolution. Trends Ecol. Evol. ISSN 0169–5347.

Westra, S., Fowler, H. J., Evans, J. P., Alexander, L. V., Berg, P., Johnson, F., Kendon, E. J., Lenderink, G. and Roberts, N. M. (2014). Future changes to the intensity and frequency of short-duration extreme rainfall. Rev. Geophys. 52, 522–555.

Whitlock, M. C. (2015). Modern approaches to local adaptation. Am. Nat. 186, S1–S4.

Wickham., H. (2016). ggplot2: Elegant Graphics for Data Analysis. Springer-Verlag *New York*.

Wolf, J. B. and Wade, M. J. (2009). What are maternal effects (and what are they not)? Philos. Trans. R. Soc. Lond. B Biol. Sci. 364, 1107–1115.

Youngson, N. A. and Whitelaw, E. (2008). Transgenerational epigenetic effects. Annu. Rev. Genomics Hum. Genet. 9, 233–257.

Zemeckis, D. R., Dean, M. J. and Cadrin, S. X. (2014). Spawning dynamics and associated management implications for Atlantic Cod. N. Am. J. Fish. Manag. 34, 424–442.

