## Supplementary Methods for "Chromosomal inversions assist acute salinity and temperature adaptation in Atlantic cod (*Gadus Morhua*) eggs"

### 1 Collection of batch 3 and 4

In batch 3, eggs were collected as stated, but the water used in the bucket was water from the laboratory instead of from the surface of the pond due to poor pond surface water conditions (low salinity). In batch 4, collection proceeded as normal; however, not enough individuals were collected during the first collection. Therefore, a second collection took place 75 minutes after the first collection. The wind was blowing on the surface water towards the north-east corner of the pond on the day of collection, and many eggs were observed floating in this corner. Five hours and 45 minutes after the 1st collection, we collected individuals (3rd collection) blown into the corner of the pond by scooping surface water with a 1 L container. We placed these individuals in a holding container, separated from batch 4 and termed them batch 4b. We used batch 4b individuals for experimentation because they reflected natural variation in the pond. The individuals that were collected from the plankton trawl were deep enough under the surface not to been blown into the corner of the pond. Furthermore, the blowing of individuals into the corner was due to the controlled habitat of the pond. In the wild, strong winds would create waves, causing the eggs to be overturned deeper in the water.

### 2 Thermal gradient block setup

We used two aluminum thermal gradient blocks to control temperatures in the experiment (Veenhof et al., 2024; Fig. 3). The 1st (A), 4th (D), and 7th (H) columns were used in the 1st block, and the 1st (K) column was used in the second block (Fig. 3). In each column, only the first 4 rows were used, except in replicates 9-13, when the 5th row was used in column K (due to a beaker getting stuck in row 4) and in replicate 13, when the 5th row was used in column H. The beaker row location was randomly determined. Tubing on each end of the blocks was connected to a Pondteam Superjet 1000 water pump placed in one of two 30 L coolers filled up to 3/4 with water. The pump with tubing connected near row A was placed in a cooler with warm water, and the pump with tubing connected near rows J and K were placed in a cooler with cold water (Fig. 3). To warm the water, a 300-watt Aqua El comfort zone water heater was placed in the "warm" cooler, and the outlet was connected to an INKBIRD Temperature Controller ITC-308 Series (Fig. 3). The probe of the temperature controller was connected to the top of the thermal gradient block, directly next to well A3 and set to heating at 17°C. When the probe temperature dropped below 17 °C, the heater was turned on until the temperature reached 17 °C again. The cold water ran through a Teco TK 6000 chiller set to 2°C via a recirculating system located in an open area and connected to an Eheim 1262210 water pump in the cooler (Fig. 3). This setup created an average (±SE) temperature of 15.9 (±0.134)°C in column A, 12.0 (±0.040)°C in column D, 7.0 (±0.025)°C in column H, and 3.3 (±0.033)°C in column K for all replicates and treatments at a given temperature.

**3 Assignment of inversion genotypes and ecotypes**

Our data was compared to a genetic baseline dataset that included 466 cod from the east Atlantic offshore, coastal, and eastern Baltic cod ecotypes, genotyped using the same SNP panel (Henriksson et al., in prep.; *cf.* Andersson et al., 2023). The baseline data set and our data set were first filtered to only include loci found in both data sets (3462 total loci). Then, any individuals for whom we did not receive SNP data from were removed, and finally, the two data sets were manually combined in a text editor (hereafter "combined data"). Afterwards, the loci in the cod locus metadata and the combined data were ordered, and loci not included in the combined data were removed from the cod locus metadata. The genomic dataset with just the eggs was tested for quality by calculating call rates and heterozygosity. The call rates for both loci and individuals were high (90.134% min call rate for loci, 95.07% min call rate for individuals), so no individuals or loci were removed. The mean and median heterozygosity were 0.34 and 0.35, respectively, determined by using the gl.report.heterozygosity function in the dartR package (v2.9.7; Gruber et al., 2018; Fig. S7). Contamination was determined to be unlikely because no extremities in heterozygosity was seen.

To assign inversion genotypes on chromosomes 2, 7, and 12, Principal Component Analyses (PCAs) were used. First, PCAs were created using the baseline data set with loci found within inv2 (18 loci), inv7 (74 loci), and inv12 (101 loci), using the dudi.pca function in the ade4 package (v1.7.22; Dray and Dufour, 2007; Fig. l). After, the eggs were plotted, one at a time to account for kinship, on the baseline dataset PCAs. To make sure individuals were plotted in the correct group, the PCA axes were oriented in the same direction each time, based on the relative locations of four individuals in the baseline dataset. Inversion genotypes for eggs were assigned based on their location relative to the baseline data set inversion groups (Fig. S8).

The individuals in the baseline dataset also acted as reference data for ecotype assignment. We used the assign.X function in the assignPOP package (v1.3.0; Chen, 2024) to calculate the percent likelihood that each individual in both the egg and baseline data sets belonged to each ecotype. We did not classify individuals into set ecotypes (ex., offshore, hybrid, and coastal) because there was no clear division of percent likelihoods that suggested divisions between ecotypes. Therefore, we used the percent likelihood of belonging to a given ecotype as our metric of interest.
