## Supplementary Figures and Tables for "Chromosomal inversions assist acute salinity and temperature adaptation in Atlantic cod (*Gadus Morhua*) eggs"

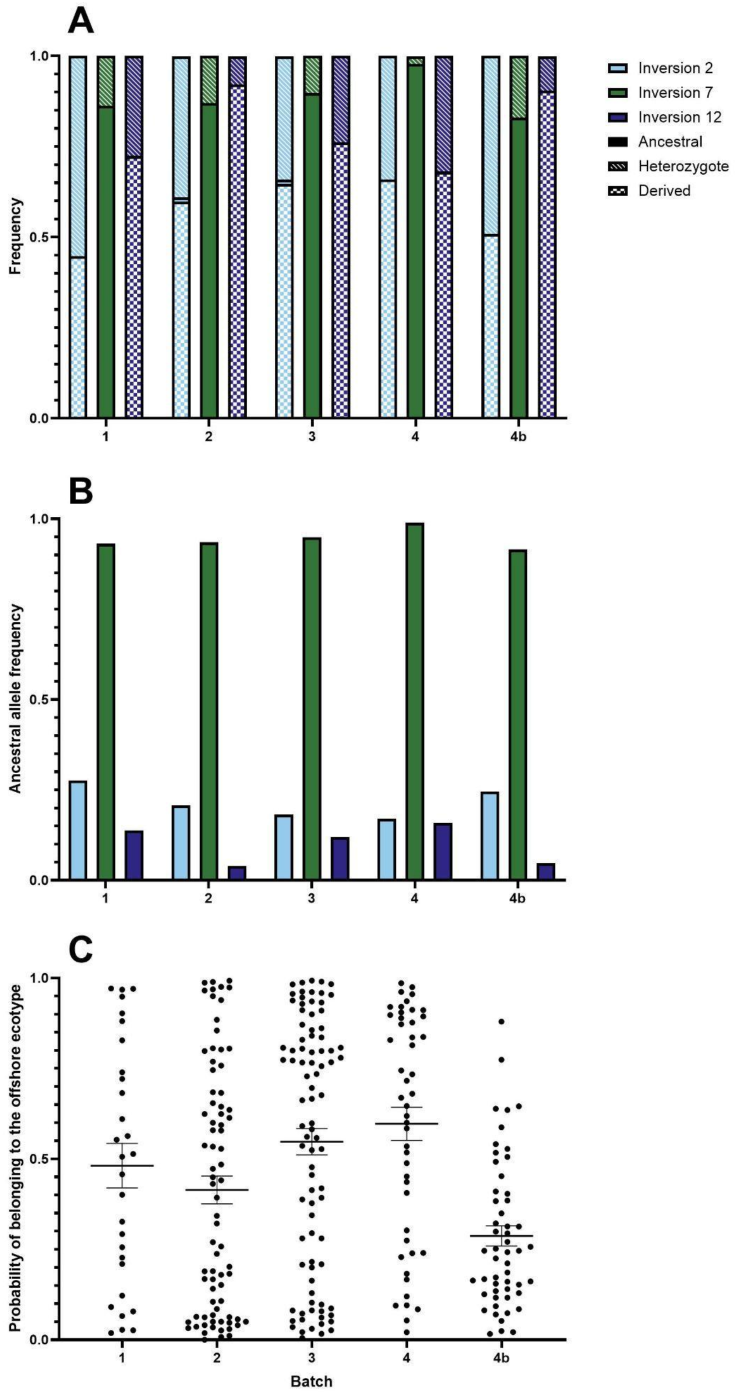


**Fig. S1.** The frequency of inversion genotypes (A) and ancestral allele (B) for inv2 (blue), inv7 (red), and inv12 (green) and the mean (±standard error) probability of belonging to the offshore ecotype (C) across batches 1, 2, 3, 4, and 4b. Genotypes include derived (solid), ancestral (checkered), and heterozygotes (angled stripes). n=30, 77, 88, 57, and 62 for batch 1, 2, 3, 4, and 4b, respectively.

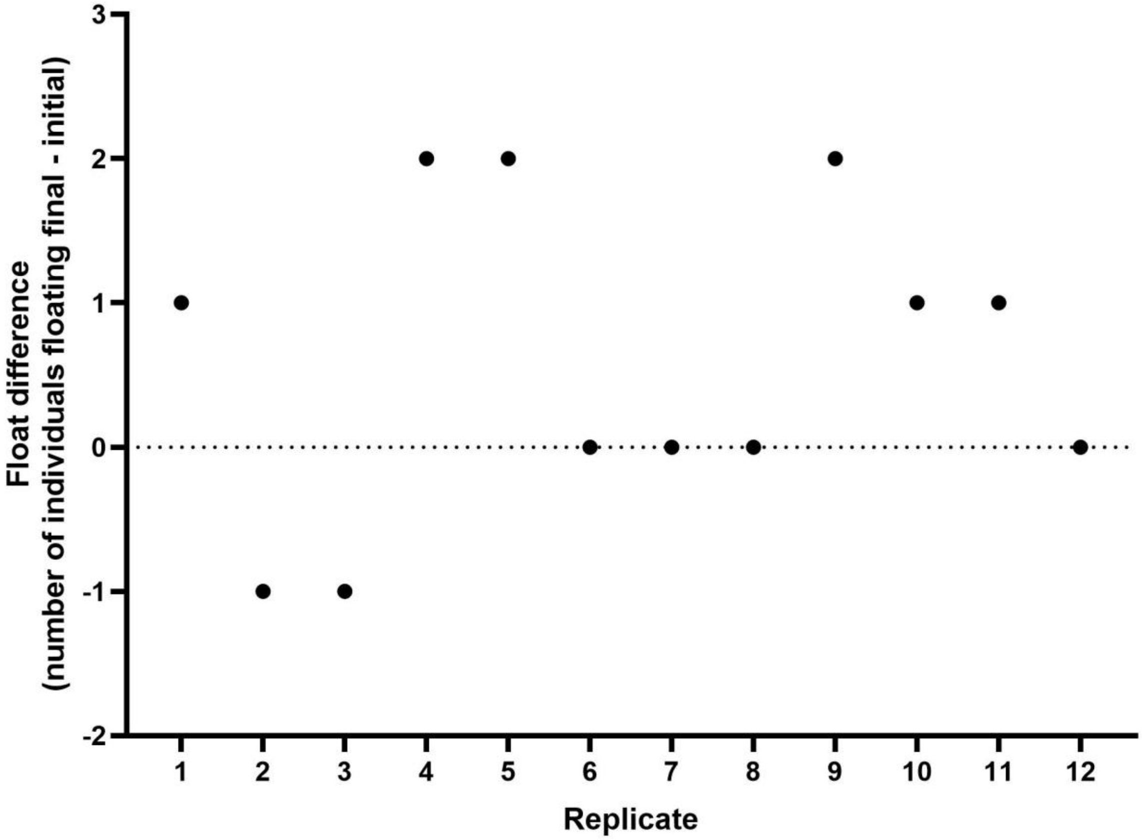


**Fig. S2.** The float difference for each replicate of the T07S30 treatment. Note how there are replicates with both positive and negative float differences, indicating that both sinking and recovery were possible.

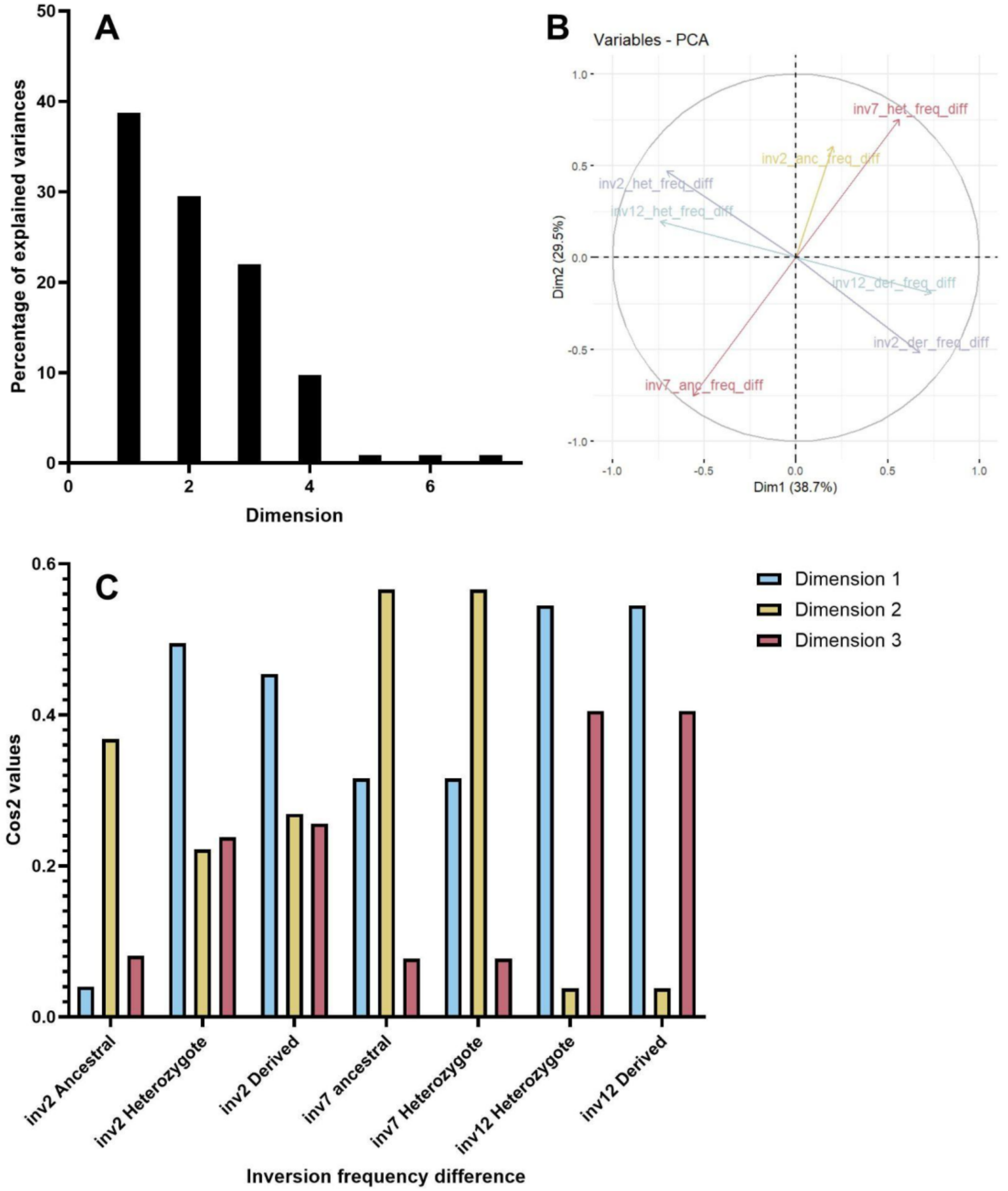


**Fig. S3.** The results from a created PCA using the inversion genotype frequency differences, including the percentage of explained variances for each dimension (A), the location of variables on the PCA coloured by their cos2 values (B), and the cos2 values for each variable along dimensions, 1 (light blue), 2 (yellow), and 3 (pink; C).

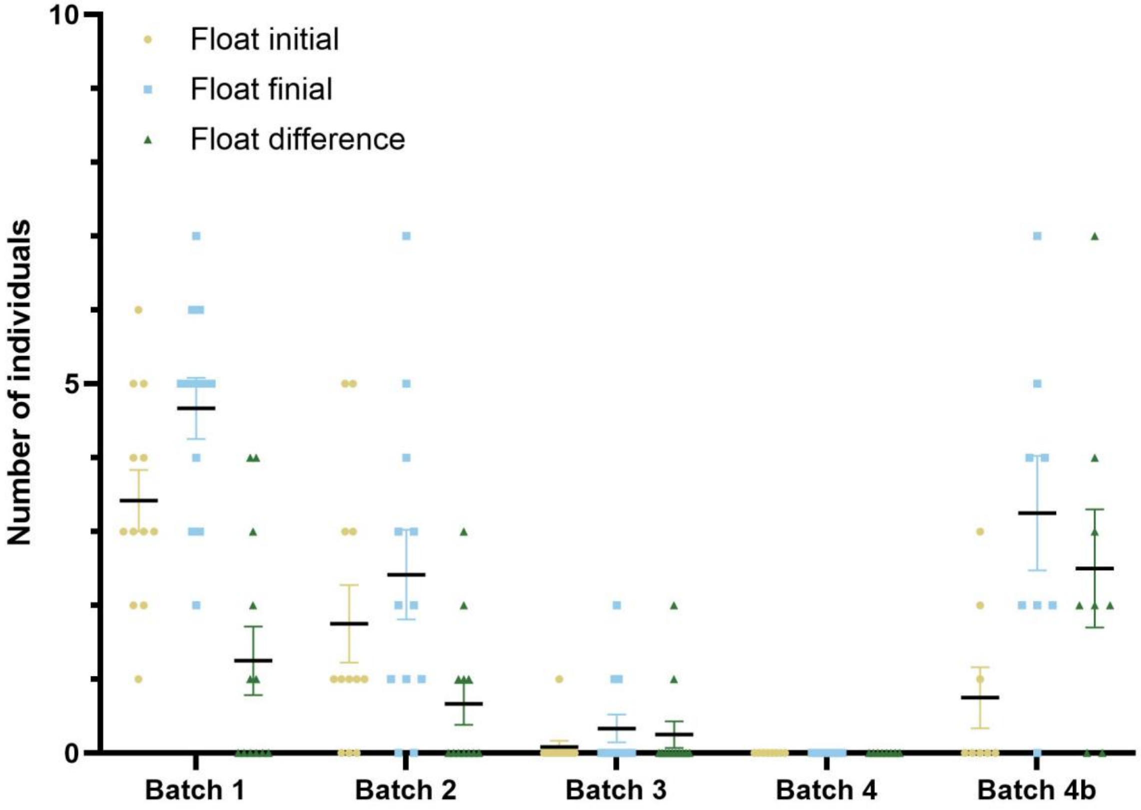


**Fig. S4.** The mean number of individuals floating initially (yellow circles) and finally (blue squares) and the float difference (green triangles) for all salinity 25 treatments vary across batches. Error bars represent the standard error. N=12, 12, 12, 8, and 8 for batches 1, 2, 3, 4, and 4b, respectively.

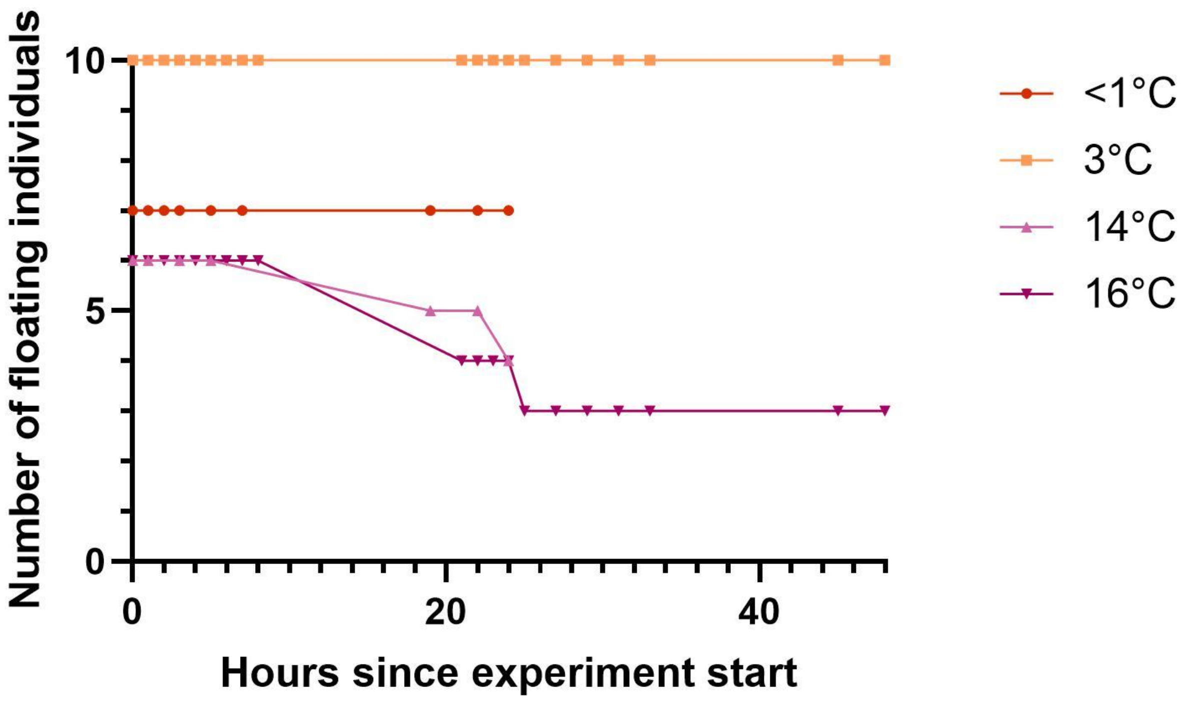


**Fig. S5.** The number of floating individuals counted at four temperatures, <1°C (n=7; red circles), 3°C (n=10; orange squares), 14°C (n=6; light purple triangles), and 16°C (n=6; dark purple inverted triangles), over 48 hours of pilot experimentation.

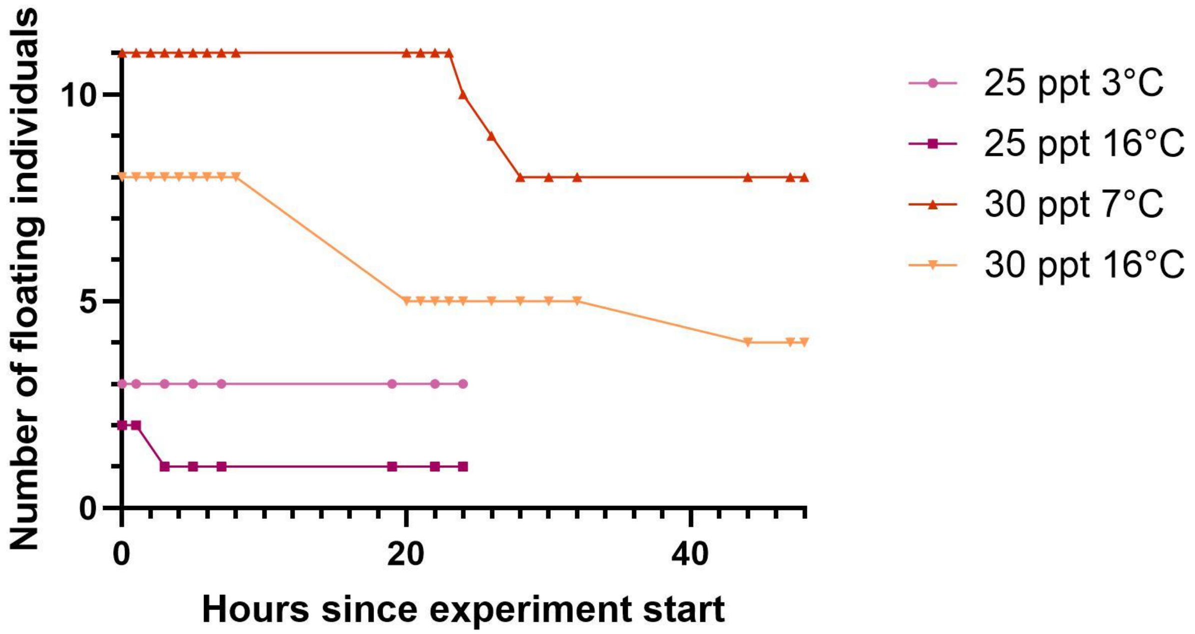


**Fig. S6.** The number of floating individuals counted in four treatments, 25 ppt at 3°C (n=8; light purple circles), 25 ppt at 16°C (n=7; dark purple squares), 35 ppt at 7°C (n=11; red triangles), and 35 ppt at 16°C (n=9; orange inverted triangles), over 48 hours of pilot experimentation.

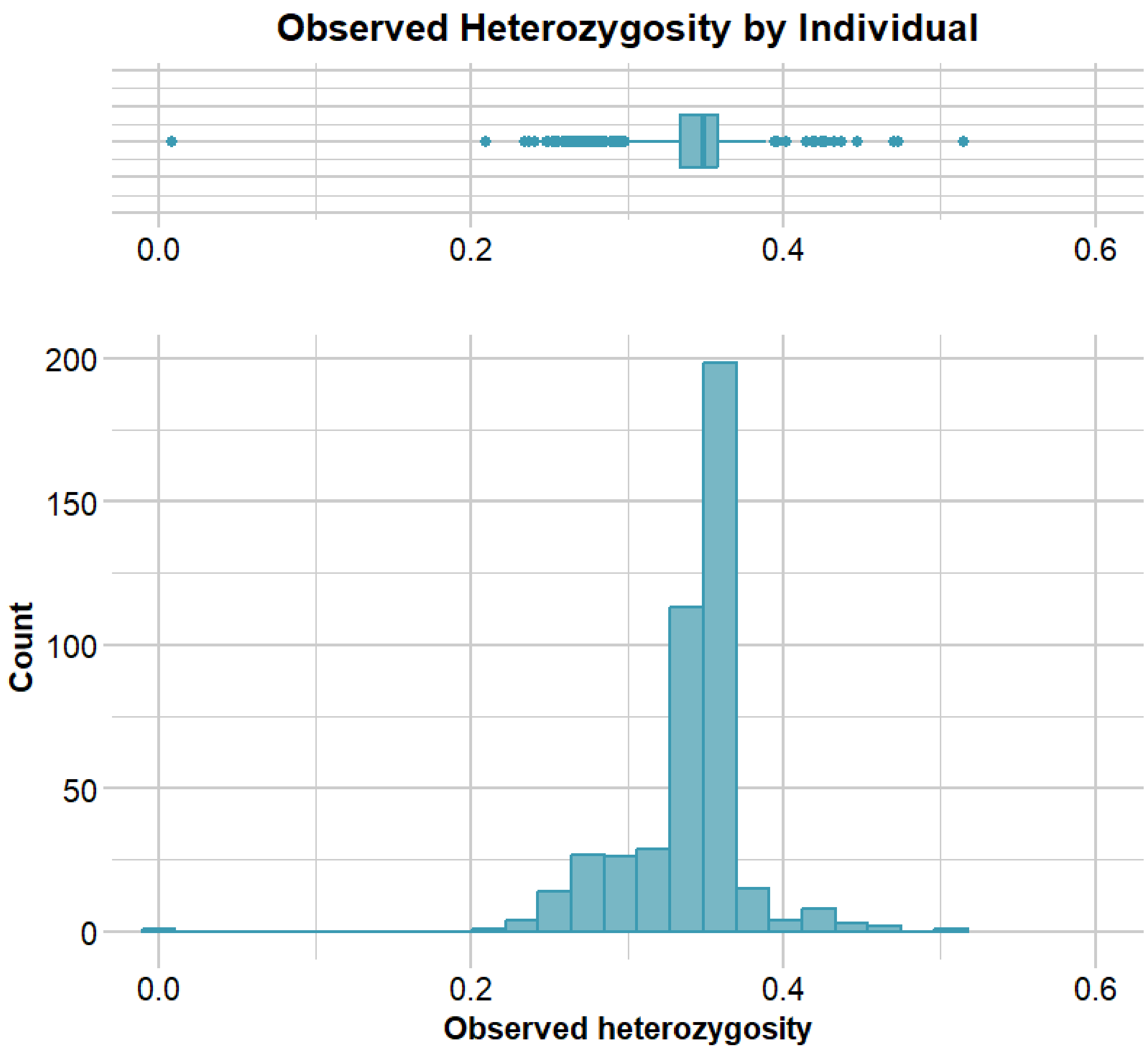


**Fig. S7.** A histogram and box plot showing the observed heterozygosity for all 446 eggs sent off for DNA analysis.

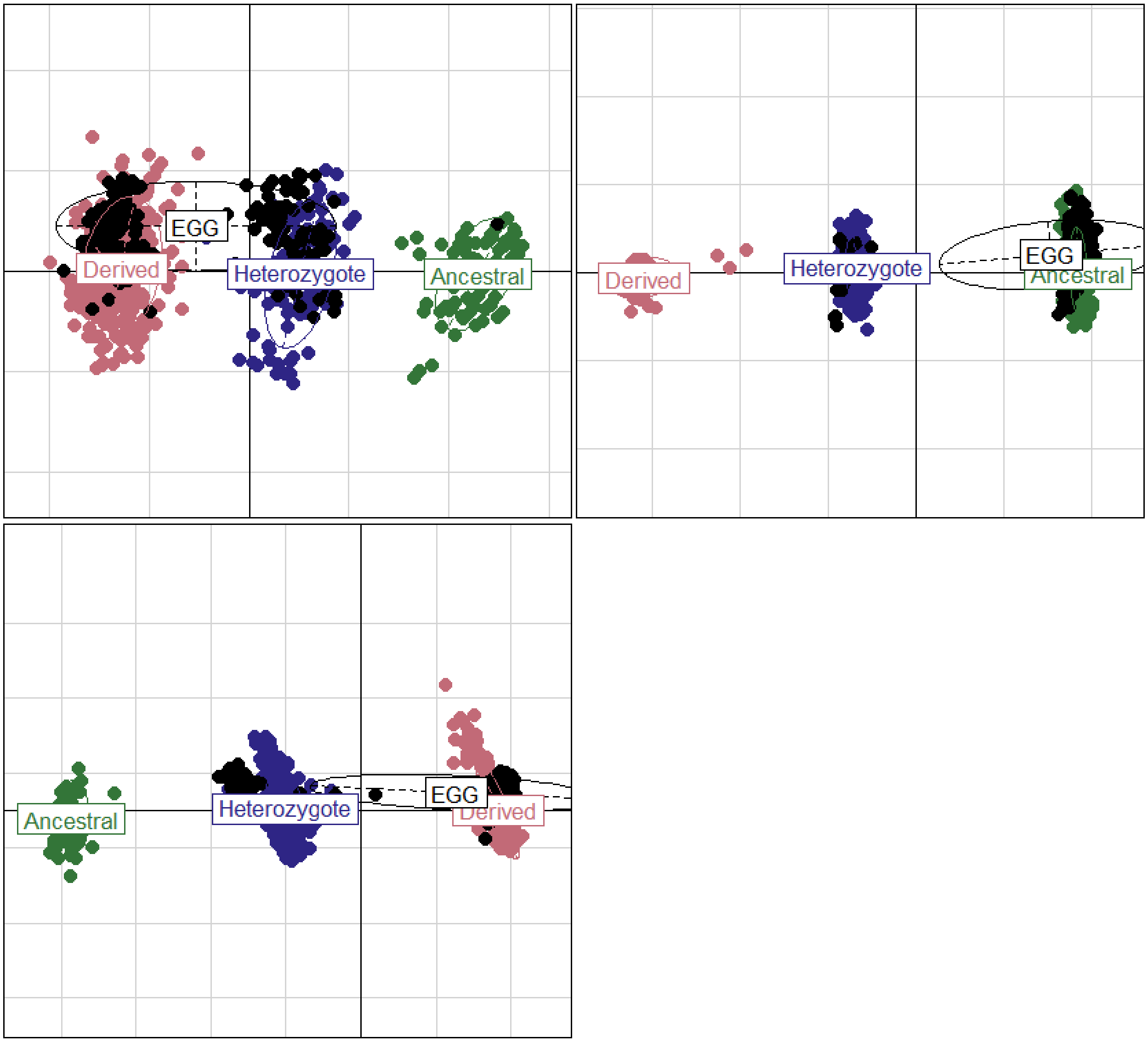


**Fig. S8.** PCA showing the genotype classifications for inv2 (A), inv7 (B), and inv12 (C) for all the individuals from the baseline dataset and where the individuals in the combined dataset are located on the PCAs.

**Table S1.** The number of individuals used in each treatment and batch of the temperature salinity stressor experiment, along with the total number of individuals per batch and treatment.

| | **1** | **2** | **3** | **4** | **4b** | **Total** |
|---|---|---|---|---|---|---|
| **T03-S25** | 30 | 30 | 30 | 20 | 20 | 130 |
| **T03-S35** | 30 | 30 | 30 | 20 | 20 | 130 |
| **T07-S20** | 30 | 30 | 30 | 20 | 10 | 120 |
| **T07-S25** | 30 | 30 | 30 | 20 | 20 | 130 |
| **T07-S30** | 30 | 30 | 29 | 19 | 10 | 118 |
| **T07-S35** | 30 | 30 | 30 | 20 | 10 | 120 |
| **T12-S25** | 30 | 30 | 30 | 20 | 20 | 130 |
| **T12-S35** | 30 | 29 | 30 | 19 | 20 | 128 |
| **T16-S25** | 29 | 30 | 30 | 20 | 20 | 129 |
| **T16-S35** | 30 | 30 | 30 | 20 | 20 | 130 |
| **Total** | 299 | 299 | 299 | 198 | 170 | 1265 |

**Table S2.** The change between the initial and final stages and states that were used to classify stage changes.

| **Classification** | **Initial stage** | **Initial state** | **Final stage** | **Final state** |
|---|---|---|---|---|
| **Early floated** | Early | Sunk | Early | Floated |
| **Sunk developed** | Early | Sunk | Late | Sunk |
| **Developed floated** | Early | Sunk | Late | Float |
| **Float developed** | Early | Float | Late | Float |
| **Late floated** | Late | Sunk | Late | Float |
| **Early sank** | Early | Float | Early | Sank |
| **Developed** | Early | Float | Late | Float |
| **Sank** | Early or Late | Float | Late | Sunk |

**Table S3.** The results from a general linear mixed model (Equation 2), including the z-value and p-value, testing the impact of temperature on five different stage changes possible in the salinity 25 ppt treatments. df=46.

| Stage change | z-value | p-value |
|---|---|---|
| early floated | -1.483 | 0.138 |
| sunk developed | 4.526 | **6.01e-6** |
| developed floated | 1.509 | 0.131 |
| float developed | 1.245 | 0.213 |
| late floated | 0.0 | 1.0 |

Bolded p-values indicate significance.

**Table S4.** The results from a general linear mixed model (Equation 2), including the z-value and p-value, testing the impact of temperature on three different stage changes possible in the salinity 35 ppt treatments. df=45.

| Stage change | z-value | p-value |
|---|---|---|
| early sank | NA | NA |
| developed | 4.315 | **1.59e-5** |
| sank | 0.5 | 0.617 |

Bolded p-values indicate significance.

**Table S5.** The results of Kolmogorov-Smirnov tests, including the d-value and p-value, comparing the probability of belonging to the offshore ecotype distributions between a batch and the four treatments, T03-S25, T07-S25, T16-S25, and T16-S35, within the batch.

| Treatment | Batch | D-value | p-value |
|---|---|---|---|
| T03-S25 | 1 | 0.1444 | 0.9531 |
| | 2 | 0.26753 | 0.8137 |
| | 3 | 0.76136 | 0.4944 |
| | 4 | NA | NA |
| | 4b | 0.56604 | 0.1223 |
| T07-S25 | 1 | 0.15047 | 0.9756 |
| | 2 | 0.28671 | 0.2678 |
| | 3 | NA | NA |
| | 4 | NA | NA |
| | 4b | 0.45283 | 0.1648 |
| T16-S25 | 1 | 0.14807 | 0.9328 |
| | 2 | 0.40909 | 0.2313 |
| | 3 | 0.72727 | 0.5618 |
| | 4 | NA | NA |
| | 4b | 0.34591 | 0.2472 |
| T16-S35 | 1 | 0.029374 | 1 |
| | 2 | 0.15634 | 0.6431 |
| | 3 | 0.075758 | 0.9994 |
| | 4 | 0.16896 | 0.7647 |
| | 4b | 0.14796 | 0.8516 |
